# Mutations in TMEM43 cause autosomal dominant auditory neuropathy spectrum disorder via interaction with connexin-mediated passive conductance channels

**DOI:** 10.1101/2020.07.27.222323

**Authors:** Minwoo Wendy Jang, Doo-Yi Oh, Eunyoung Yi, Xuezhong Liu, Jie Ling, Nayoung Kim, Kushal Sharma, Tai Young Kim, Seungmin Lee, Ah-Reum Kim, Min Young Kim, Min-A Kim, Mingyu Lee, Jin-Hee Han, Jae Joon Han, Hye-Rim Park, Bong Jik Kim, Sang-Yeon Lee, Dong Ho Woo, Jayoung Oh, Soo-Jin Oh, Tingting Du, Ja-Won Koo, Seung Ha Oh, Hyun Woo Shin, Moon-Woo Seong, Kyu Yup Lee, Un-Kyung Kim, Jung Bum Shin, Shushan Sang, Xinzhang Cai, Lingyun Mei, Chufeng He, Susan H. Blanton, Zheng-Yi Chen, Hongsheng Chen, Xianlin Liu, Aida Nourbakhsh, Zaohua Huang, Woong-Yang Park, Yong Feng, C. Justin Lee, Byung Yoon Choi

**Affiliations:** KU-KIST Graduate School of Converging Science and Technology, Korea University, Seoul 02841, Korea; Center for Cognition and Sociality, Institute for Basic Science (IBS), Daejeon 34141, Korea; Department of Otorhinolaryngology, Seoul National University Bundang Hospital, Seongnam 13620, Korea; College of Pharmacy and Natural Medicine Research Institute, Mokpo National University, Mokpo, Korea; Department of Otolaryngology, University of Miami Miller School of Medicine, Miami, FL, USA; Dr. John T. Macdonald Foundation Department of Human Genetics and Hussman Institute for Human Genomics, University of Miami Miller School of Medicine, Miami, FL, USA; Department of Otolaryngology, Xiangya Hospital, Central South University, Changsha, Hunan, China; Institute of Molecular Precision Medicine, Xiangya Hospital, Central South University, Changsha, Hunan, China; Samsung Genome Institute, Samsung Medical Center, Seoul, Korea; Department of Biology, College of Natural Sciences, Kyungpook National University, Daegu, 41566, Republic of Korea; School of Life Sciences, KNU Creative BioResearch Group (BK21 plus project), Kyungpook National University, Daegu, 41566, Republic of Korea; Department of Biomedical Sciences, Seoul National University Graduate School, Seoul, Korea; Department of Otorhinolaryngology, Seoul National University Hospital, Seoul, Korea; Research Center for Animal Model, Jeonbuk Department of Inhalation Research, Korea Institute of Toxicology, KRICT, Jeongeup 56212, Korea; Convergence Research Center for Diagnosis, Treatment and Care System of Dementia, Korea Institute of Science and Technology (KIST) Seoul 02792, Korea; Department of Neuroscience, University of Virginia, Charlottesville, Virginia 22908, USA; Department of Laboratory Medicine, Seoul National University Hospital, Seoul National University College of Medicine, Seoul, Korea; Department of Otorhinolaryngology-Head and Neck Surgery, Kyungpook National University Hospital, Kyungpook National University School of Medicine, Daegu, Korea; Department of Otology and Laryngology, Harvard Medical School and Eaton-Peabody Laboratory, Massachusetts Eye and Ear Infirmary, Boston 02114, USA; Changsha Central Hospital, University of South China, Changsha, Hunan, China

## Abstract

Genes that are primarily expressed in cochlear glia-like supporting cells (GLSs) have never been clearly associated with progressive deafness. Herein, we present a novel deafness locus mapped to chromosome 3p25.1 and a new auditory neuropathy spectrum disorder (ANSD) gene *TMEM43* mainly expressed in GLSs. We identify p.R372X of *TMEM43* by linkage analysis and exome sequencing in two large Asian families. The knock-in (KI) mouse with p.R372X mutation recapitulates a progressive hearing loss with histological abnormalities exclusively in GLSs. Mechanistically, TMEM43 interacts with Cx26 and Cx30 gap junction channels, disrupting the passive conductance current in GLSs in a dominant-negative fashion when the p.R372X mutation is introduced. Based on the mechanistic insights, cochlear implant was performed on two patients and speech discrimination was successfully restored. Our study highlights a pathological role of cochlear GLSs by identifying a novel deafness gene and its causal relationship with ANSD.

---

The main organ for hearing and speech discrimination is the organ of Corti of inner ear, and its major cell types include neuron-like inner hair cells (IHCs) and outer hair cells (OHCs) and glia-like supporting cells (GLSs). GLSs have been suggested to play an important role in development and maintenance of auditory system^1^. These cells are defined as glia-like cells due to typical glia markers such as GFAP and GLAST^2^. Mutations of connexin (Cx) channels of GLSs such as Cx26 (GJB2)^3,4^ and Cx30 (GJB6)^4,5^ are known to cause hearing impairment, highlighting the importance of GLSs in deafness. However, the underlying molecular mechanisms of how these gap junction channels interact with other proteins and lead to hearing loss still remain to be explored.

Hearing loss is defined as a diminished sensitivity to the sounds normally heard^6^ or an inability in speech discrimination despite preserved sensitivity to sound as in auditory neuropathy spectrum disorder (ANSD). As of 2017, about 1.4 billion people worldwide are affected by hearing loss^7^ The causes of hearing loss include exposure to aging, ototoxic drugs, other environmental insults, and most importantly, genetic alterations which account for 60% of the hearing loss (http://hereditaryhearingloss.org). Recent advances in sequencing technologies have facilitated discovery of a genetic etiology of deafness with varying degrees and features^8,9^ Especially, understanding the roles and functions of novel genes in the auditory system has been made through a stepwise approach: cloning of a novel deafness gene and investigation of the function by *in vitro* studies and analyses on mutant animal models^10–12^. Over one hundred deafness genes have been identified in hair cells (http://hereditaryhearingloss.org). However, the genes in GLSs and their roles in hearing or speech discrimination, especially their contribution to ANSD still remains unknown.

## Results

### *TMEM43* is a novel deafness gene

We identified two large five-generation pedigrees of Chinese Han family (HN66) and Korean family (SB162), non-consanguineously segregating the adult-onset progressive ANSD in an autosomal dominant fashion (**Fig. 1a** and **Extended Data Fig. 1a**). Unlike the normal subject (#290) in SB162, the affected subjects (#17, #291, #284, #304) from families HN66 and SB162 displayed elevated pure tone audiogram (PTA) thresholds and disproportionately lower speech discrimination score (SDS) for the PTA thresholds. Importantly, complete absence of auditory brainstem response (ABR) was noted from these affected subjects, despite the presence of either distortion-product otoacoustic emission (DPOAE) or cochlear microphonics (CM) (**Fig. 1b**), which are classic signs of ANSD. CM measurement was not performed from Subjects #17 and #36. These typical symptoms did not begin to appear in a full extent until the age of thirties. The youngest subject (#36 from HN66, a 16-year-old female) showed a significant elevation of ABR threshold as high as 80dB, while DPOAE response of #36 was completely preserved (**Fig. 1b**). This suggests that phenotype for #36 can be interpreted as the stage of transition to canonical ANSD.

**Figure 1.**
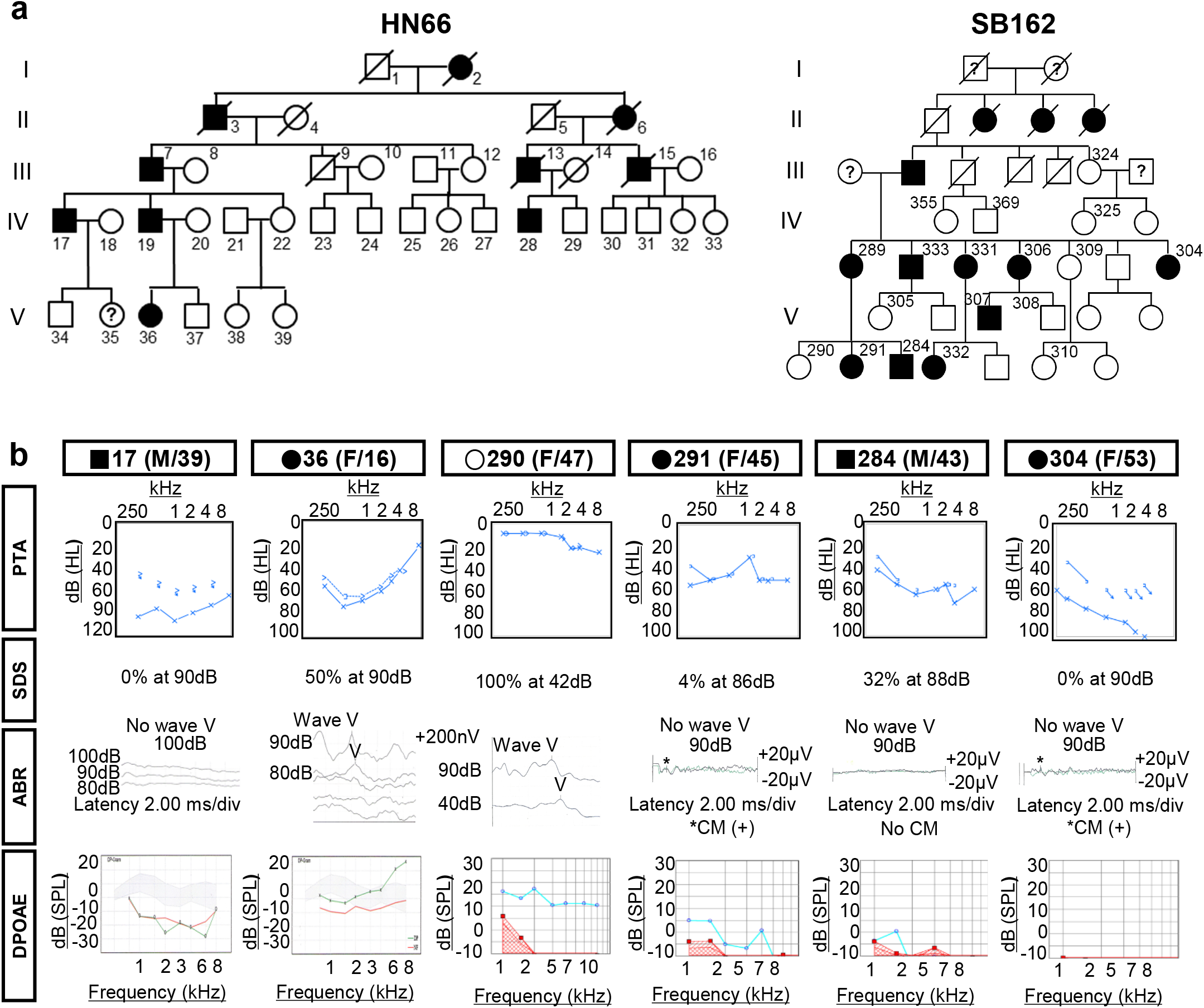
Pedigree and audiological assessment of family HN66 and SB162. **a**, A Chinese family HN66 and a Korean family SB162 segregates ANSD in an autosomal dominant fashion. Open and filled symbols indicate unaffected and affected statuses, respectively. Square and circle symbols indicate male and female respectively. Subject numbers are superscripted. **b**, PTA, SDS, ABR and DPOAE of 6 subjects from HN66 and SB162. Unaffected subject #290 shows a definite normal ABR response to sound stimulus as low as 40dB. Despite absence of ABR response, affected subjects (#17, #291, #284, #304) show either presence of DPOAE response (blue line) or cochlear microphonics (CM) (asterisk), indicating ANSD from these subjects. The youngest subject (#36 from HN66, a 16-year-old female) among all affected members from the two families showed obviously disordered waveform differentiation in ABR responses and significant elevation of ABR threshold up to as high as 80dB, while DPOAE response of #36 was completely preserved. This suggests that phenotype for #36 can be interpreted as the stage of transition to canonical ANSD.

Stepwise genetic analysis was carried out to identify the pathogenic variants of families HN66 and SB162, separately (**Fig. 2a**). To identify the chromosomal locus of ANSD in the two families, we conducted a whole-genome linkage-scan on 9 subjects (5 affected and 4 unaffected) in HN66 and 18 subjects (10 affected and 8 unaffected) in SB162 (**Extended Data Fig. 1c**). As a result, we identified a candidate region on chromosome 3: 13,165,401-22,769,511 with the highest parametric log_10_ *Odds* (LOD) value of 2.4 across the entire chromosomes in HN66 (**Fig. 2b, left**) and two regions (Region #1: Chr 3: 1,946,000 – 5,956,000; Region #2: Chr 3:11,883,000-14,502,000) in the chromosome 3p25.1 with a genetic length of 6.7 and 3 cM, each displaying a significant LOD score greater than 3.0, respectively, in SB162 (**Fig. 2b, right** and **Extended Data Fig. 2a inset**). Given the failure to show any link with the previously reported ANSD genes (*OTOF* (NM_194248, 2p23.3)^13^, *DIAPH3* (NM_001042517, 13q21.2)^14^, *SLC17A8* (NM_139319, 12q23.1)^15^, *AIFM1* (NM_004208, Xq26.1)^16^, and *OPA1* (NM_130837, 3q29)^17^) and a new ANSD gene reported by us, *ATP1A3* (NM_152296.4, 19q13.2)^18^, we suspected an involvement of a novel gene.

**Figure 2.**
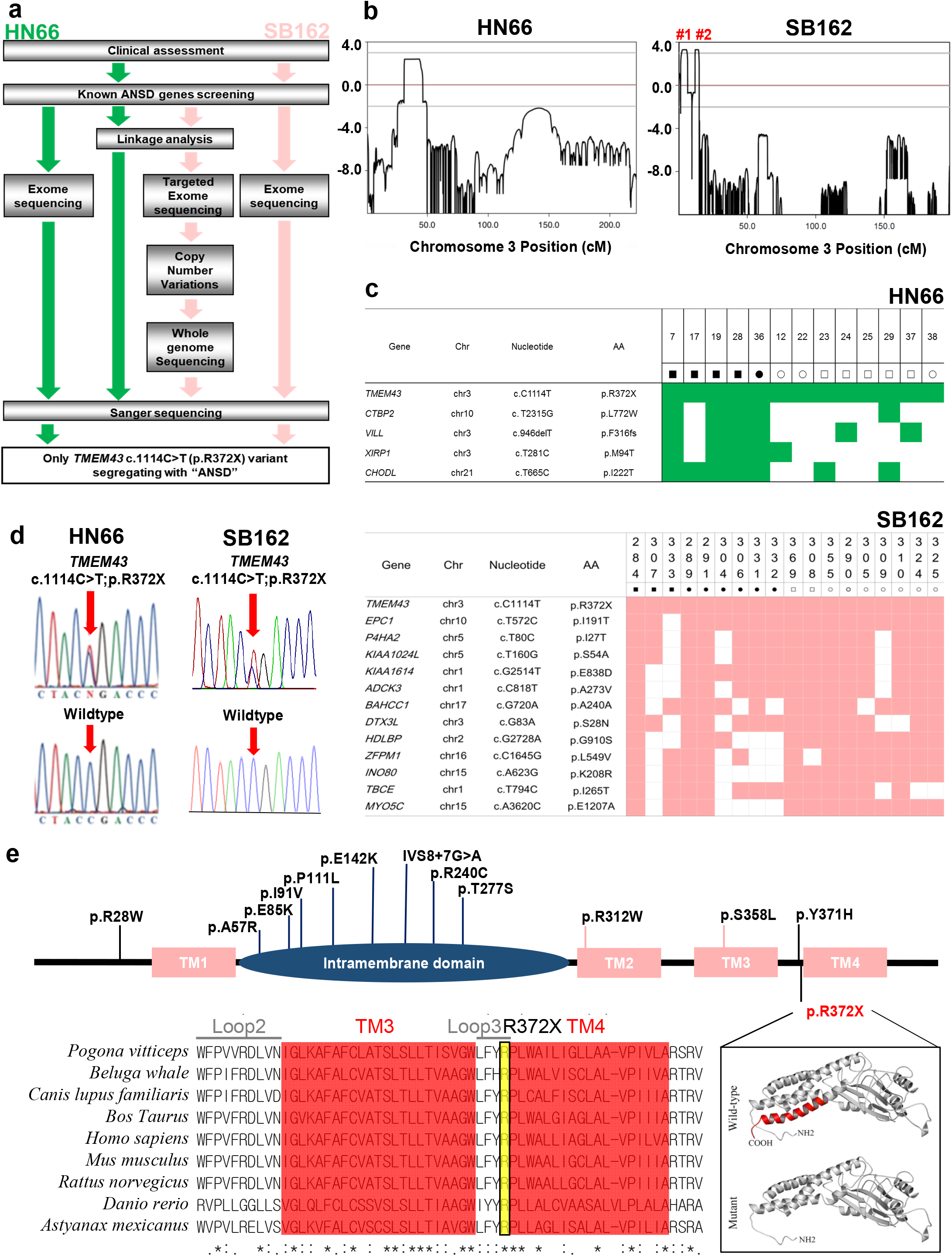
A p.R372X variant of *TMEM43* is the causative variant of hearing loss in family HN66 and SB162. **a**, Overall workflow of variant detecting in families HN66 and SB162. **b**, Merlin multipoint linkage analysis demonstrates LOD scores greater than 3.0 only on chromosome 3p25.1. Region #1 (Chr. 3: 1,946,000 – 5,956,000), region #2 (Chr. 3:11,883,000-14,502,000). **c**, Genetic candidates of the ANSD after exome sequencing of genomic DNA from 3 subjects (2 affected subjects (#36 and #28) and 1 unaffected subject (#22)) in HN66 and 4 subjects (the 3 affected (#289, #291, #284) and 1 unaffected, (#290)) in SB162 are listed. Subsequent segregation study with additional subjects to identify *TMEM43* in HN66 and SB162 as the only remaining candidate with full-match. Co-segregation of a gene variant with the hearing loss phenotype is indicated as green and an apricot colored box, while mismatch is shown as a white box. Chr; chromosome, AA; amino acid. **d**, DNA sequence chromatograms of c.1114C>T variant in *TMEM43*. **e**, The mutation spectrum of TMEM43 associated with ARVD5 and EDMD7 reported in HGMD database (Top, black letter) and the variants identified in the present study (Bottom, p.R372X). The conservative prediction of the amino acid sequences of TMEM43 is shown below. Asterisks indicate the amino acids that are identical across all species. Three-dimensional modeling of wild-type human TMEM43 and the variant TMEM43, lacking TM4, identified in the present study. TM, transmembrane domain.

Although the identification of a novel causative gene for ANSD from these two families was done independently and blindly in China and South Korea, the linkage interval shared by the two families narrowed down to Chr 3: 13,165,401 −14,502,000. Exome sequencing was first performed from two affected subjects (#36 and #28) and one unaffected subject (#22), yielding five candidate variants co-segregating with the phenotype (*XIRP1, CTBP2, CHODL, TMEM43* and *VILL*) (**Supplementary Table 1**). Next, we performed detailed Sanger sequencing of these variants from 10 additional subjects in HN66, leaving only one variant (p.R372X from *TMEM43*) which co-segregated with the ANSD phenotype (**Fig. 2c,d**). This variant fell within the linkage the linkage interval of HN66 (chr3: 13,165,401-22,769,511). For SB162, we employed two diagnostic pipelines in a parallel fashion **(Fig. 2a)**. First, we performed targeted exome-sequencing of the two regions obtained from the linkage analysis among four subjects (#284, #289, #290, #291), leading to identification of two nonsynonymous variants (*TMEM43*-p.R372X and *FBLA2*-p.D851H) in Region #2 that co-segregated with the ANSD phenotype (**Table 1, top**). Of the two variants, we excluded *FBLA2*-p.D851H, based on its high occurrence (0.4%) among the healthy Korean population in two independent databases, KRGDB (n=1722) and in-house Samsung Genome Institute control subjects database (n=400), leaving p.R372X from *TMEM43* (NM_024334) as the only candidate. No convincing copy number variations, such as a large genomic deletion and duplication, were detected within the locus (**Extended Data Fig. 2b**). To exclude non-coding causative variants, we also performed whole genome sequencing of the linkage interval from eight subjects (#284, #289, #290, #304, #307, #309, #324, #332). The filtering step left us one nonsynonymous variant (TMEM43-p.R372X) and eighteen non-coding region variants that co-segregated with the deaf phenotype within the linkage interval (**Supplementary Table 2**). We excluded all of the eighteen variants, based on their high occurrence (>0.4%) among the two aforementioned Korean databases from normal controls and detection of these variants among unaffected subjects or normal controls (n=65) by Sanger sequencing, again leaving p.R372X from TMEM43 (NM_024334) as the only candidate (**Supplementary Table 2**). Separately from the first pipeline and independently of linkage analysis, we performed exome sequencing from the same four subjects (#284, #289, #290, #291), identifying 20 possible candidate variants through an extensive filtering process in SB162 (**Table 1, bottom**). We performed detailed Sanger sequencing of these variants from 14 additional subjects in the SB162, again leaving only one variant (p.R372X from *TMEM43*) as a candidate (**Fig. 2c,d** and **Extended Data Fig. 1b**).

**Table 1.** Nonsynonymous variants within the locus* co-segregating with ANSD that survived filtering process after exome sequencing among 4 subjects from SB162.

| Func | Gene | ExonicFunc | GenbankID | Position | Exon | Nucleotide | AA | Chr | Start | End | Ref | Alt | Global MAF | KRGDB<br>(N=1722) | PolyPhen-2 | SIFT | GERF++ |
| --- | --- | --- | --- | --- | --- | --- | --- | --- | --- | --- | --- | --- | --- | --- | --- | --- | --- |
| **Targeted exome sequencing after linkage analysis** |  |  |  |  |  |  |  |  |  |  |  |  |  |  |  |  |  |
| exonic | TMEM43 | Stopgain SNV | NM_024334 | Region #<br>2<sup>**</sup><br>(3p25.1) | exon12 | c.C1114T | p.R372X | chr3 | 14,183,206 | 14,183,206 | C | T | T=0.000008/1 (ExAC) | ND | NA | NA | 4.83 |
| exonic | FBLN2 | nonsynonymous SNV | NM_001998 | Region #<br>2<sup>**</sup><br>(3p25.1) | exon11 | c.G2551C | p.D851H | chr3 | 13,670,527 | 13,670,527 | G | C | C=0.0007/84 (ExAC)<br>C=0.0004/2 (1000 Genomes) | C=0.005517/19 | 0.893 | 0.04 | 4.23 |
| **Exome sequencing** |  |  |  |  |  |  |  |  |  |  |  |  |  |  |  |  |  |
| exonic | KIAA1614 | nonsynonymous SNV | NM_020950 |  | exon5 | c.G2514T | p.E838D | chr1 | 180905559 | 180905559 | G | T | ND | ND | 0.002 | 0.39 | -5.05 |
| exonic | ADCK3 | nonsynonymous SNV | NM_020247 |  | exon6 | c.C818T | p.A273V | chr1 | 227169815 | 227169815 | C | T | T=0.00006/5 (ExAC)<br>T=0.00008/1 (GO-ESP)<br>T=0.0001/4 (TOPMED) | ND | 0.999 | 0.04 | 5.1 |
| exonic | TBCE | nonsynonymous SNV | NM_001079515 |  | exon9 | c.T794C | p.I265T | chr1 | 235599116 | 235599116 | T | C | ND | ND | 0.01 | 0.62 | 4.61 |
| exonic | PARD3B | nonsynonymous SNV | NM_057177 |  | exon11 | c.A1594C | p.I532L | chr2 | 206023605 | 206023605 | A | C | ND | ND | 0.28 | 0.08 | -1.97 |
| exonic | HDLBP | nonsynonymous SNV | NM_001243900 |  | exon21 | c.G2728A | p.G910S | chr2 | 242176107 | 242176107 | C | T | T=0.000008/1 (ExAC)<br>T=0.0002/1 (1000 Genomes) | ND | 0.094 | 0.64 | 0.666 |
| exonic | TMEM43 | stopgain SNV | NM_024334 |  | exon12 | c.C1114T | p.R372X | chr3 | 14183206 | 14183206 | C | T | T=0.000008/1 (ExAC) | ND | NA | NA | 4.83 |
| exonic | DTX3L | nonsynonymous SNV | NM_138287 |  | exon1 | c.G83A | p.S28N | chr3 | 122283356 | 122283356 | G | A | ND | ND | 0.454 | 0.03 | -9.39 |
| exonic | KIAA1024L | nonsynonymous SNV | NM_001257308 |  | exon1 | c.T160G | p.S54A | chr5 | 129084043 | 129084043 | T | G | ND | ND | NA | NA | 3.53 |
| exonic | P4HA2 | nonsynonymous SNV | NM_001017973 |  | exon2 | c.T80C | p.I27T | chr5 | 131554240 | 131554240 | A | G | ND | G=0.00029/1 | 0.01 | 0.07 | 6.17 |
| exonic | DOK3 | nonsynonymous SNV | NM_001144875 |  | exon2 | c.C8T | p.P3L | chr5 | 176936534 | 176936534 | G | A | ND | ND | 0.991 | 0.03 | 4.47 |
| exonic | MAML1 | nonsynonymous SNV | NM_014757 |  | exon5 | c.C2338T | p.H780Y | chr5 | 179201165 | 179201165 | C | T | ND | T=0.00029/1 | 0.916 | 0.31 | 4.94 |
| exonic | ARID1B | nonframeshift insertion | NM_017519 |  | exon1, exon1 | c.1029_1030insGCG | p.A343delinsAA | chr6 | 157100092 | 157100092 | - | GC<br>G | ND | InsGCG=0.001742/6 | NA | NA | -0.657 |
| exonic | EPC1 | nonsynonymous SNV | NM_001272004 |  | exon4 | c.T572C | p.I191T | chr10 | 32582010 | 32582010 | A | G | ND | ND | 0.999 | 0.07 | 6.04 |
| exonic | CRYAB | nonsynonymous SNV | NM_001885 |  | exon1 | c.T79A | p.F27I | chr11 | 111782370 | 111782370 | A | T | ND | ND | 0.002 | 0.35 | 2.56 |
| exonic | INO80 | nonsynonymous SNV | NM_017553 |  | exon6 | c.A623G | p.K208R | chr15 | 41379795 | 41379795 | T | C | C=0.0001/17 (ExAC) | C=0.0046457/16 | 0.009 | 0.61 | 4.2 |
| exonic | MYO5C | nonsynonymous SNV | NM_018728 |  | exon29 | c.A3620C | p.E1207A | chr15 | 52515748 | 52515748 | T | G | ND | ND | 0.365 | 0.73 | 5.97 |
| exonic | CILP | nonsynonymous SNV | NM_003613 |  | exon5 | c.C448G | p.R150G | chr15 | 65497781 | 65497781 | G | C | ND | ND | 0.0 | 0.67 | 0.901 |
| exonic | MAP2K1 | nonsynonymous SNV | NM_002755 |  | exon10 | c.A1062C | p.Q354H | chr15 | 66782095 | 66782095 | A | C | C=0.00002/2 (ExAC) | C=0.001161/4 | 0.001 | 0.02 | -5.39 |
| exonic | ZFPM1 | nonsynonymous SNV | NM_153813 |  | exon10 | c.C1645G | p.L549V | chr16 | 88600011 | 88600011 | C | G | ND | G=0.00029/1 | 0.978 | 0.01 | 2.61 |
| exonic | MIDN | nonsynonymous SNV | NM_177401 |  | exon5 | c.C401T | p.S134L | chr19 | 1254182 | 1254182 | C | T | T=0.00003/2 (ExAC)<br>T=0.0002/5 (TOPMED) | ND | 1 | 0 | 3.8 |

<sup>\*</sup> Locus: Region 1+ Region 2 (See Figure 1c inset); <sup>\*\*</sup> Region #1: Chr 3: 1,946,000 - 5,956,000; Region #2: Chr 3:11,883,000-14,502,000; Ref: Reference sequence; Alt: Alternate sequence; ND: not detected; NA: not applicable.
The candidate variants were firstly listed up if they were nonsynonymous variants in coding regions. Variants with a minor allele frequency less than 1% were then selected based on the Exome Sequencing Project 6500 (ESP6500), the 1000 Genome Project (1000G), The Exome Aggregation Consortium (ExAC), Trans-Omics for Precision Medicine (TOPMED) and our in-house database containing the exomes of 81 Korean individuals. Following the inheritance pattern, homozygous variants and compound heterozygote variants with sufficient read depths (>10X) and a genotype quality (>20) were selected commonly found in the affected siblings. Finally, the variants which have none of clinical significance of dbSNP ID were finally identified. In silico prediction Algorithm: Polyphen-2 (http://genetics.bwh.harvard.edu/pph2/index.shtml) ; SIFT (http://sift.jcvi.org/www/SIFT_chr_coords_submit.html); Conservation tools: GERP++ score in the UCSC Genome Browser (http://genome-asia.ucsc.edu/) ; ExAC, Exome Aggregation Consortium (http://exac.broadinstitute.org/) ; 1000 Genomes (https://www.ncbi.nlm.nih.gov/variation/tools/1000genomes/) ; KRGDB, Korean Reference Genome DB (http://152.99.75.168/KRGDB/); TOPMED, Trans-Omics for Precision Medicine (https://www.nhlbiwgs.org/).

Along with the mutation spectrum of *TMEM43* associated with the heart diseases reported in HGMD database (**Fig. 2e, top, black letter**), the arginine at position 372, as well as amino acids in the fourth transmembrane domain (TM) and C-terminal of TMEM43, is highly conserved across organisms (**Fig. 2e, bottom**). A 3D modeling of human *TMEM43* showed that p.R372X, which was expected to introduce a premature stop codon, caused a truncation of the last 29 amino acids of the TMEM43 protein. (**Fig. 2e, inset**). This variant was predicted to be pathogenic, according to the guideline for the interpretation of classifying pathogenic variants (PS3, PM2, PP1_Strong and PS4_Supporting)^19,20^ and high CADD score of 45 (https://cadd.gs.washington.edu/info)^21^. Previously, p.S358L in *TMEM43* has been shown to cause familial arrhythmogenic right ventricular cardiomyopathy (ARVC)^22,23^. However, our ANSD subject of HN66 and SB162 displayed symptoms of neither arrhythmia nor any other heart abnormalities. Conversely, hearing loss has not been reported in *TMEM43-p.S358L* ARVC patients^22,23^. This suggests that p.S358L and p.R372X of *TMEM43* exert a completely different pathophysiological mechanism, leading to pleiotropy of this gene.

### TMEM43 is expressed in GLSs

The cochlear expression of TMEM43 was firstly confirmed by RT-PCR (**Fig. 3a**). TMEM43 was expressed throughout the mouse body including cochlea, heart, eye and brain (**Fig. 3a**). To specify TMEM43 expression in the cochlea tissue, we performed immunostaining using anti-TMEM43 antibody with epitope targeting p100-p200 (Ab_1-1_). The expression of TMEM43 protein was evaluated at various time points along the postnatal development of hearing in the mouse cochlea (**Fig. 3b-d**). The previous studies have reported that TMEM43 is localized predominantly to the inner nuclear membrane in multiple non-cardiac cell types and also shows some expression outside the nucleus, including the endoplasmic reticulum^23,24^ In the case of *ex vivo* cochlea, TMEM43 was mainly expressed in the organ of Corti, along the entire cochlear length, at postnatal day 4 (P4) through P20 (**Fig. 3b-e**). At P20, the expression became more restricted to the subset of the inner supporting cells (a subtype of GLSs) of the cochlea, which was the apical membrane of the inner border cells and the cell junctions of the inner sulcus cells (**Fig. 3d**). Throughout the early developmental period, up to P20, the TMEM43 expression was found exclusively in GLSs, but not in the hair cells, at both protein and mRNA levels. (**Fig. 3b-i** and **Supplementary Video 1, 2**). Adult mice at 1, 2, and 4 months still maintained this restricted expression in GLSs (**Extended Data Fig. 3a**). This glia-specific TMEM43 expression was recapitulated in adult primates corresponding to age 40’s in humans, showing exclusive expression in the junctional area between the inner supporting and IHCs, without any overlap with calretinin staining (**Extended Data Fig. 3b**). TMEM43 expression in cochlear glial cell was confirmed again by co-expression with glia specific GFAP signal **(Extended Data Fig. 3c**). The specificity of the antibody was confirmed via a significant reduction of immunoreactivity from the cultured mouse organ of Corti when infected with lentivirus carrying *Tmem43-shRNA* (**Extended Data Fig. 4a,b**). These results strongly suggest that TMEM43 may play a critical role in cochlea GLSs.

**Figure 3.**
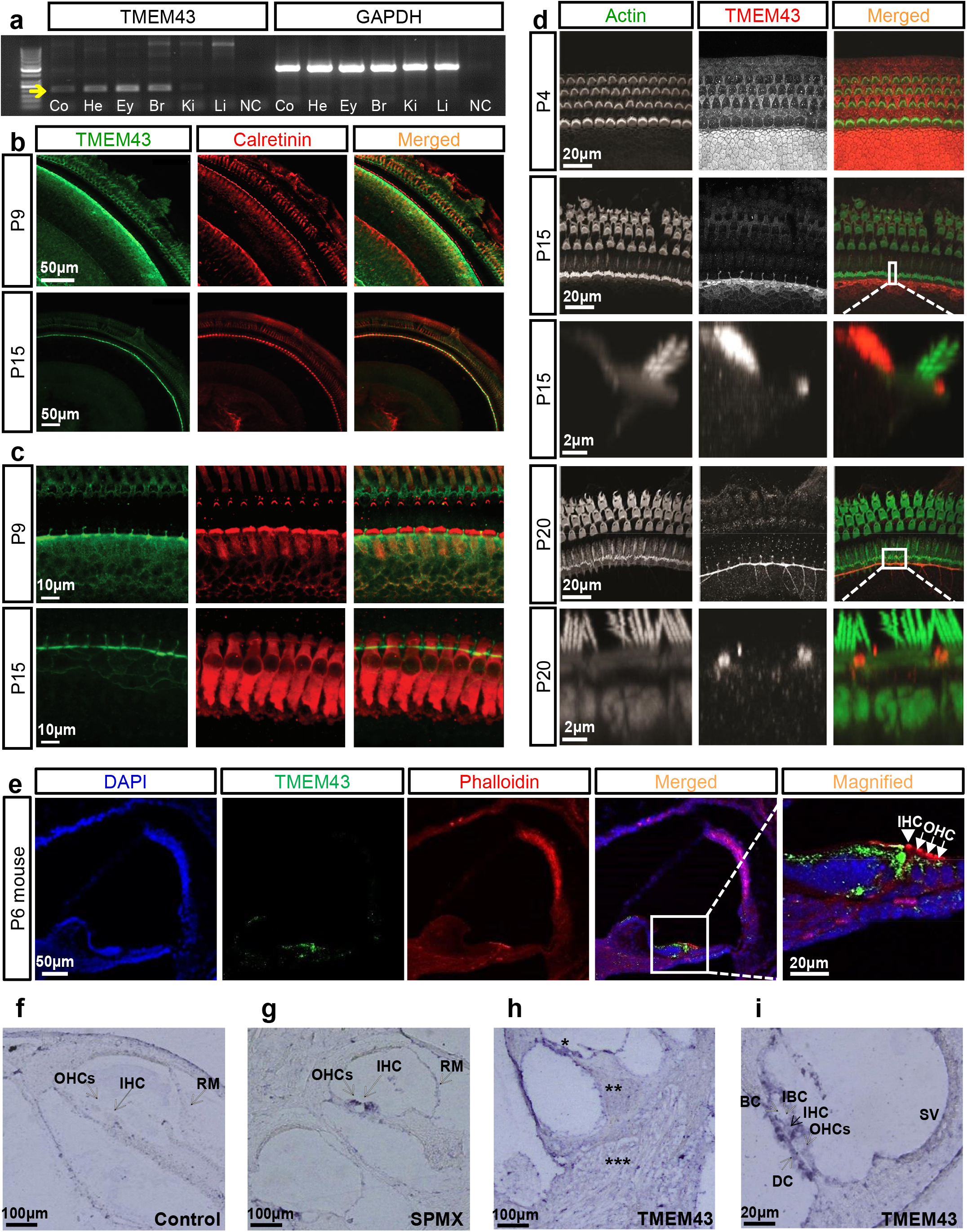
TMEM43 is exclusively expressed in the GLSs of mouse cochlea. **a**, RT-PCR gel analysis of TMEM43 in various regions of mouse. Arrow indicates TMEM43 expression (size 196bp) (Co, cochlea; He, heart; Ey, eye; Br, brain; Ki, kidney; Li, liver; NC, negative control). **b,c** Low magnification (**b**) and high magnification (**c**) confocal micrographs of cochlear turns with anti-TMEM43 (Ab_1-1_) (green) and anti-calretinin (red). **d**, Spatiotemporal expression of TMEM43 (red) and phalloidin (green) at the mouse organ of Corti. TMEM43 immunoreactivity detected widely in the Kolliker’s organ at P4 becomes restricted in the course of postnatal development at inner border cells and cell junctions of the inner sulcus cells at P20. Boxed area indicates magnified view of TMEM43 localization at the apical membrane or cortex of the inner border cell. **e**, Frozen-section images of the mouse cochlea at P6 immunolabeled by TMEM43 (Ab_1-1_) (green) and counterstained by phalloidin (red). TMEM43 immunoreactivity is detected mainly at the organ of Corti and the magnified image shows TMEM43 immunoreactivity exclusively at the GLSs of the organ of Corti, not in the hair cells. **f-i**, In situ hybridization with a scrambled probe (**f**), with *Tmem43* probe showing *Tmem43* mRNA expression in the Organ of corti (*) but not in the spiral ganglion neuron (**) and cochlear nerve (***) (**g**), with hair cell marker SMPX (**h**), and data with higher magnification (**i**). IHC; Inner hair cell, OHC; Outer hair cell, IBC; Inner border cell, BC; border cell, DC; Deiters cells, RM; Reisners membrane.

### Localization of TMEM43 in the plasma membrane and loss of protein stability in *TMEM43* p.R372X

Based on hydrophobicity analysis, the secondary structure of TMEM43 was predicted to have four transmembrane domains and one intramembrane domain with its N- and C-terminal domains located at extracellular space^22^ (**Extended Data Fig. 5a**). Bioinformatics revealed that TMEM43 was phylogenetically close among many species (**Extended Data Fig. 5b**) and the four transmembrane domains were highly conserved among different species (**Extended Data Fig. 5c**). We verified the predicted TMEM43 topology through immunocytochemistry with or without cell permeabilization. FLAG-tagged TMEM43 was transfected to HEK293T and was double stained with anti-FLAG and anti-TMEM43 with epitope targeting p203-p308 (Ab_1-2_), at intracellular loop1 (**Fig. 4a**). We found that TMEM43 immunoreactivity was positive only with cell permeabilization, indicating that the loop1 of TMEM43 resides at the intracellular space (**Fig. 4b**). In addition, a positive FLAG signal without cell permeabilization was detected **(Fig. 4b),** indicative of membrane expression of TMEM43 and confirming the topology. To verify the cell-surface localization of TMEM43, we performed cell surface biotinylation assay. We found that TMEM43 wildtype (WT) protein trafficked to the plasma membrane, whereas TMEM43-p.R372X showed almost complete disappearance of the biotinylated protein along with a significant reduction in total protein (**Fig. 4c**). Analysis of protein stability by the cycloheximide chase (CHX) assay showed that this reduction was due to a decreased protein stability of TMEM43-p.R372X **(Fig. 4d)**. Together, these results explain the etiology of TMEM43-p.R372X by reduction of protein expression at plasma membrane due to decreased protein stability.

**Figure 4.**
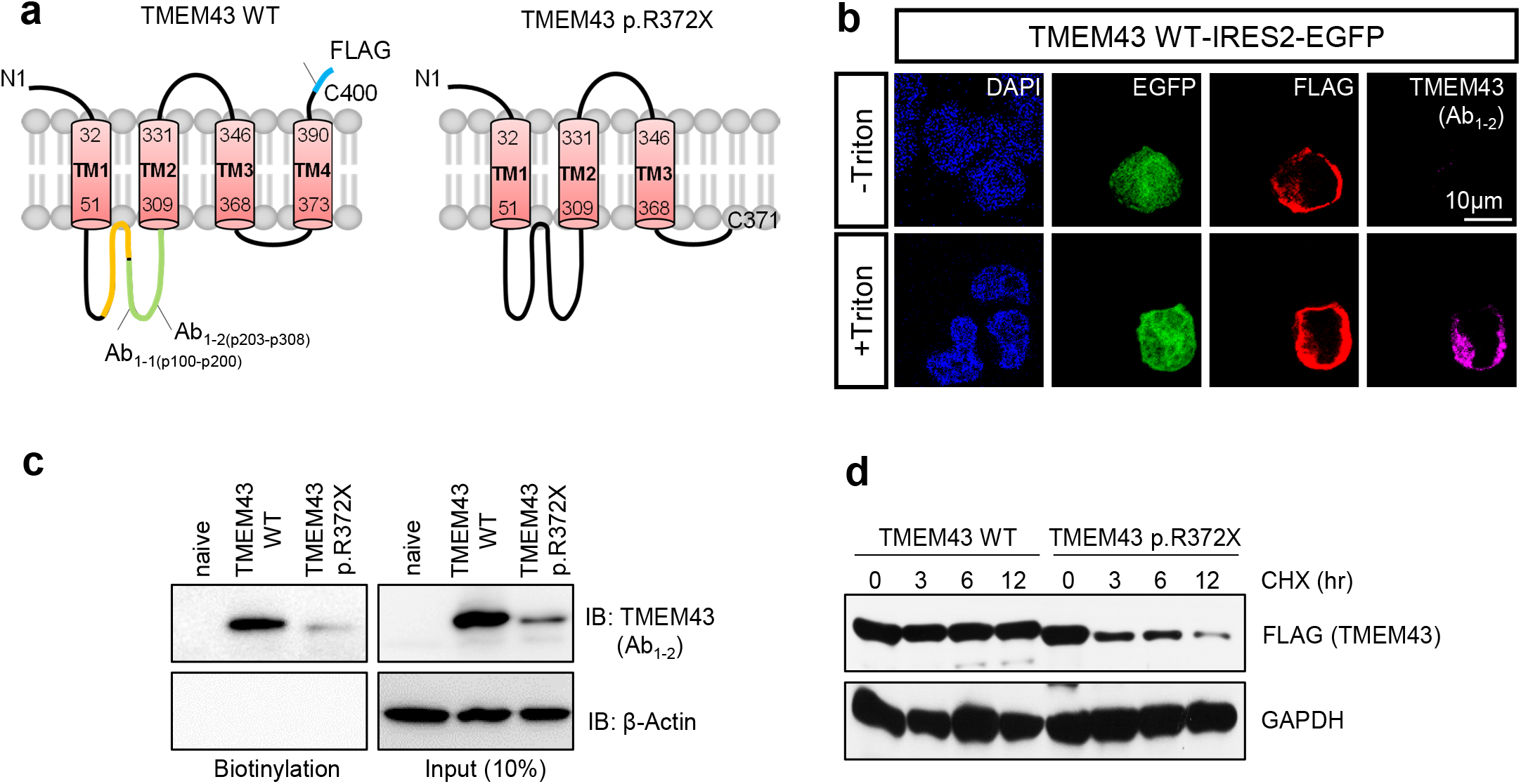
p.R372X of TMEM43 shows decreased cell surface expression with reduced protein stability. **a,** 2D topology of TMEM43 WT and p.R372X. Antibody epitopes are indicated as Ab_1-1_ and Ab_1__2 and FLAG is tagged at C-terminal. **b,** Immunostaining result of *TMEM43* expressing HEK293T cells with and without cell permeabilization. Cells were double stained with FLAG and TMEM43 antibody (Ab_1-2_). **c**, Cell surface biotinylation assay in *TMEM43* WT and p.R372X transfected HEK293T cells blotted with TMEM43 antibody (Ab_1-2_). **d**, Cycloheximide (CHX) assay with FLAG antibody. 40 ug/ml CHX was used. Band sizes for TMEM43 WT and TMEM43-p.R372X are 44kD and 41kD each.

### Generation of p.R372X knock-in mice displaying morphological change in GLSs with progressive hearing loss

To examine the pathogenic effect of mutant TMEM43 on the *in vivo* function of cochlea, a knock-in (KI) mouse harboring the human variant, p.R372X of *TMEM43* in the corresponding mouse *Tmem43* sequence (C57BL/6J;129S-*Tmem43^tm1Cby^*) was successfully established (**Fig. 5a**). The mutant TMEM43 protein was present in the GLSs till the adult period, as shown by a positive immunostaining from *Tmem43^KI^* mouse models **(Fig. 5b, Extended Data Fig. 3d**). Scanning electron microscopic (SEM) examinations showed that the apical surface area of GLSs, specifically the inner border cells of the organ of Corti from *Tmem43^+/KI^* became significantly narrower than that of *Tmem43^+/+^* from 7 months of age corresponding to mid-thirties in humans. This reduced size of GLSs became more prominent at 13 months of age in *Tmem43^+/KI^* (**Fig. 5c,d**). In contrast, the shape and the arrangement of hair cell stereocilia as well as the number of synaptic ribbons were not different between the two genotypes (**Extended Data Fig. 6a,b**). Moreover, there was no sign of severe degeneration of OHCs up to 13 months in *Tmem43^+/KI^* mice compared with the littermate controls (*Tmem43^+/+^* mice) (**Extended Data Fig. 6c**). These results indicate that p.R372X KI mice display morphological dysfunction specifically in GLSs. Finally, we observed a tendency of elevation of ABR thresholds starting at frequencies of 16 and 32 kHz from both *Tmem43^+/KI^* and *Tmem43^KI/KI^* mice compared with littermate *Tmem43^+/+^* control mice at 6 months of age which matches the onset time of hearing loss in human patients **(Fig. 5e,f)**. Significantly reduced wave I amplitude from *Tmem43^+/KI^* and *Tmem43^KI/KI^* compared with that from *Tmem43^+/+^* was noted, even at 8 kHz at 6 months of age **(Fig. 5g)**, but not at 8 kHz at 3 months of age **(Fig. 5h)**. The DPOAE responses representing the function of outer hair cells in both *Tmem43^+/KI^* and *Tmem43^KI/KI^* mice were significantly higher than the noise level at the frequencies from 7.2 to 9.7 kHz, suggesting preservation of outer hair cell function. There was no significant difference in the sound levels of DPOAE among *Tmem43^+/+^, Tmem43^+/KI^*, and *Tmem43^KI/KI^* mice (ANOVA, p > 0.05) at whole kHz (**Extended Data Fig. 6d**). These features of KI mice reflect the characteristics of ANSD^25^, also indicating a similar pathogenic effect of the human *TMEM43* variant on mice. In contrast, the electrocardiography of the *Tmem43^+/KI^* mice did not show any sign of ARVC (**Extended Data Fig. 7a**), again confirming a profoundly distinct pathogenic effect of p.R372X compared to p.S358L.

**Figure 5.**
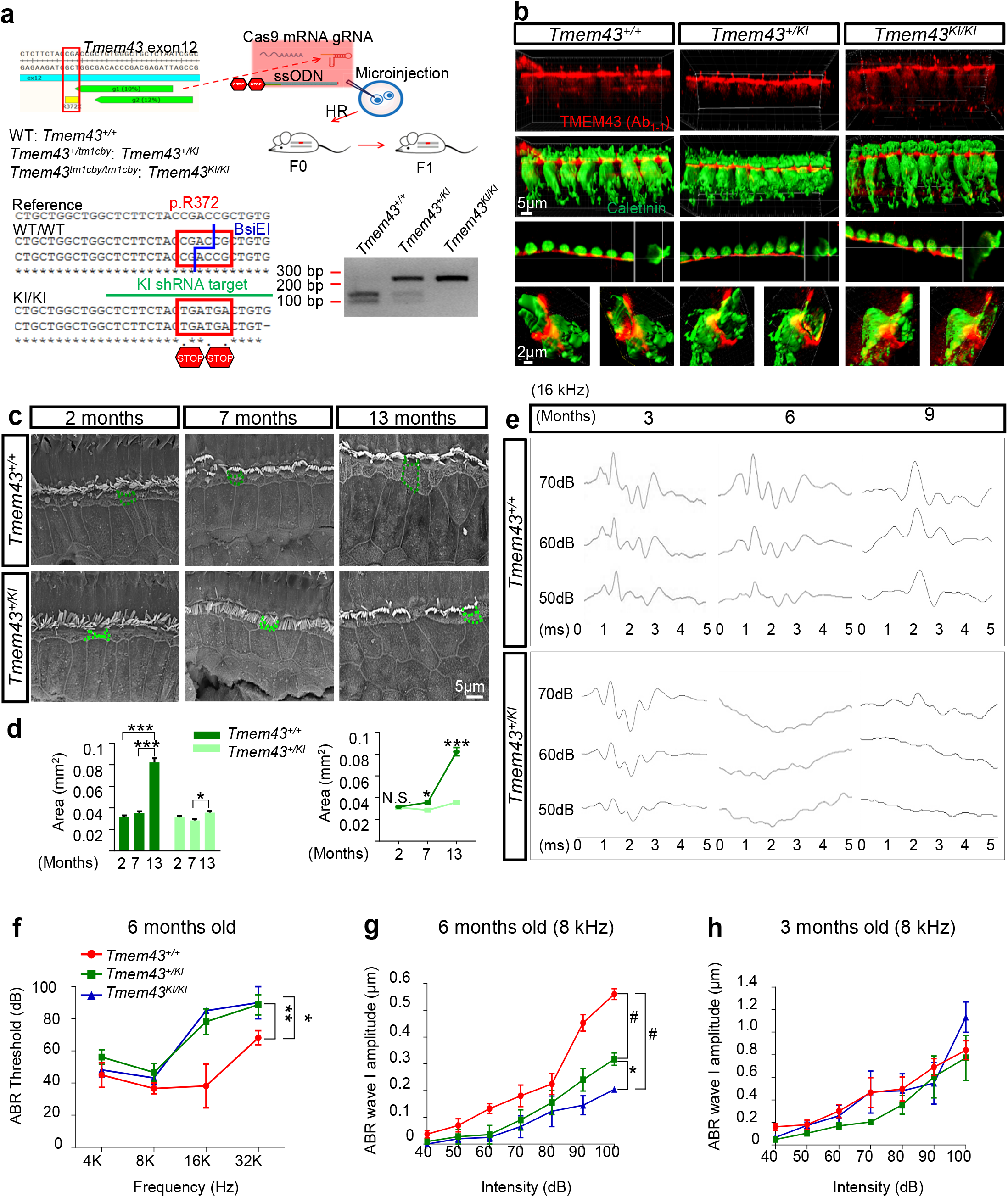
TMEM43-p.R372X KI mice display morphological and physiological dysfunction. **a**, Schematic flow of generation of *Tmem43*-p.R372X KI (C57BL/6J;129S-*Tmem43^tm1Cby^*) mice model using Clustered Regularly Interspaced Short Palindromic Repeats (CRISPR) technology. Double stop codons were inserted at p.R372. A germline transmission of p.R372X was confirmed. The p.R372X site, KI-targeting shRNA sequence and restriction enzyme site of BsiEI are each noted with red, green and blue color. Genotyping result using BsiEI is shown. gRNA: guide RNA, ssODN: Single-stranded oligodeoxynucleotide donor, HR: homologous recombination. **b**, Confocal micrographs of the organs of Corti from age-matched (1 month old) littermates of *Tmem43^+/+^, Tmem43^+/KI^* and *Tmem43^KI/KI^*. 3D reconstruction images were obtained from the apical turn of cochlea immunolabeled with anti-TMEM43 (Ab_1-1_) (red) and anti-calretinin (green). XZ and YZ section views (third row) and surface object clipping images of a single IHC (fourth row) show that TMEM43 immunoreactivity is found primarily in the apical potion of inner border cell, juxtaposing the upper lateral membrane of calretinin-positive IHC. **c**, SEM findings at the level of reticular lamina of hair cells. Green and light green dotted line each indicates apical surface area of inner border cells of *Tmem43^+/+^* and *Tmem43^+/KI^* respectively. **d**, Summary graphs of apical surface area of inner border cells (n=50) from (**c**)**. e**, Auditory brainstem response (ABR) waveforms recorded in *Tmem43^+/+^* and *Tmem43^+/KI^* mice at 3, 6, and 9 months. The x axis indicates the different time, and the y axis indicates the sound pressure level (SPL). 6 and 9 months *Tmem43^+/KI^* mice show no detectable wave response to 50 and 60dB tone burst sound in ABR, in contrast to the *Tmem43^+/+^* mice, indicative of progressive deafness. **f**, Mean ± SEM ABR thresholds from littermate *Tmem43^+/+^* (red, n=3), *Tmem43^+/KI^* (green, n=12), and *Tmem43^KI/KI^* (blue, n=3) measured at 6 months after birth. **g**, Mean ± SEM ABR wave I amplitude growth function for littermate *Tmem43^+/+^* (red, n=4), *Tmem43^+/KI^* (green, n=4) and *Tmem43^KI/KI^* (blue, n=4) at 8 kHz measured at 6 months after birth. **h,** Mean ± SEM ABR wave I amplitude growth function for *Tmem43^+/+^* (blue, n=3), *Tmem43^+/KI^* (red, n=3) and *Tmem43^KI/KI^* (green, n=3) at 8 kHz measured at 3 months after birth. ABR wave I amplitudes were not significantly different among each genotype at whole decibels measured.

### TMEM43 is an essential component of connexin channels in cochlear GLSs

TMEM43 has been previously shown to be critical for gap junction channel function in cardiac muscles^23^, raising a possibility that pathogenic effect of TMEM43-p.R372X involves gap junction channels in cochlea such as Cx26 or Cx30^26^. Therefore, we examined the localization of TMEM43 with connexin channels. Immunostaining data showed that TMEM43 was expressed in GLSs along with Cx26 and Cx30 (**Fig. 6a**). In addition, the cochlea of *Tmem43^+/+^, Tmem43^+/KI^* and *Tmem43^KI/KI^* all showed similar Cx26 expression level **(Fig. 6b)**. To directly confirm the physical interaction of TMEM43 with these connexin channels, we next performed co-immunoprecipitation assay. Both TMEM43 WT and TMEM43-p.R372X protein were immunoprecipitated with either Cx26 or Cx30 protein (**Fig. 6c**). These results together indicate that TMEM43 interact with connexin channels.

**Figure 6.**
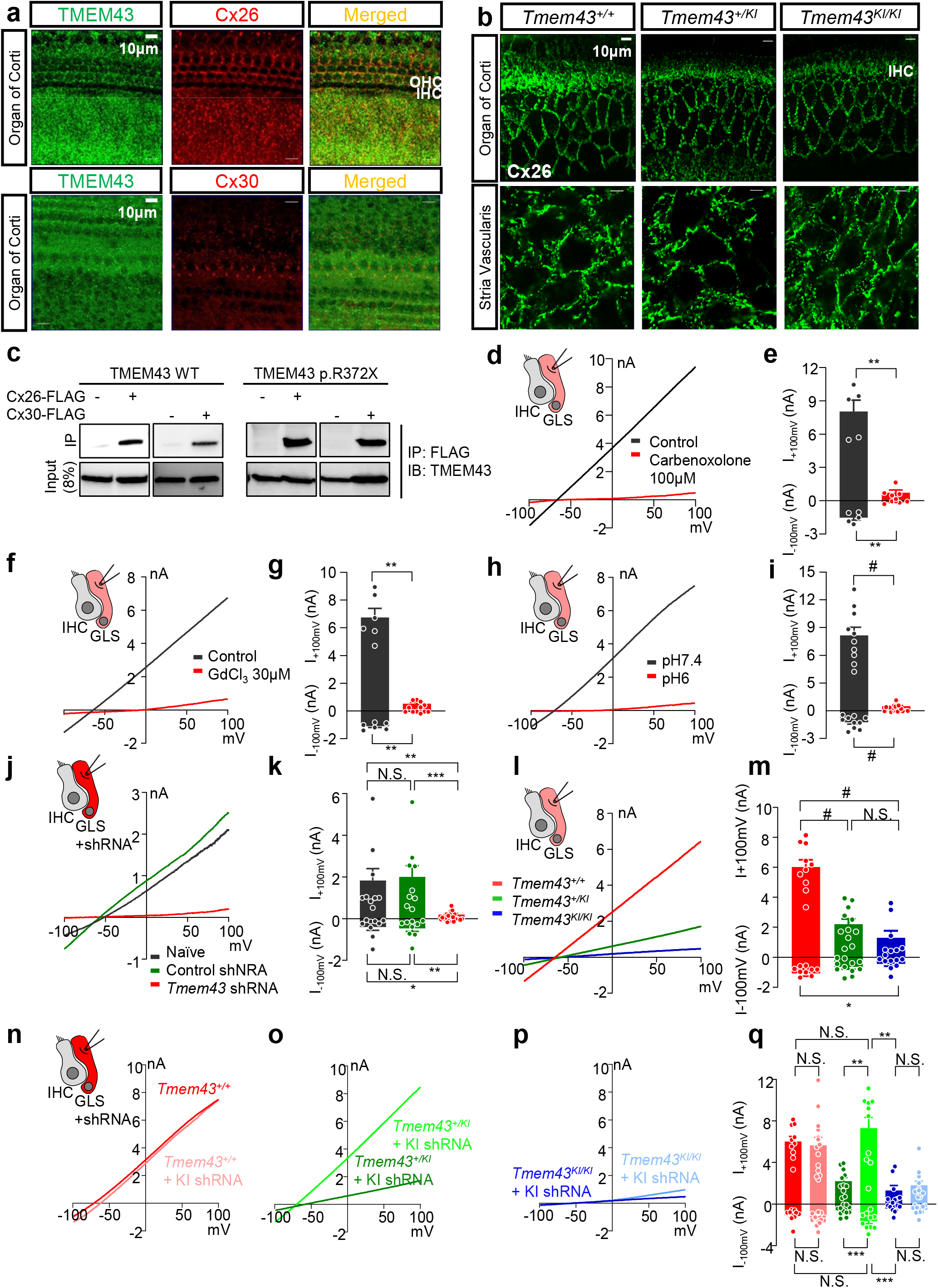
TMEM43 is an essential component of connexin channels to induce passive conductance current in GLSs. **a**, Immunohistochemistry data showing TMEM43 expression with Cx26 (upper panel), and with Cx30 (bottom panel) in GLSs (P6). **b**, Confocal micrographs of Cx26 expression in Organ of Corti (upper panel) and Stria Vascularis (bottom panel) of *Tmem43^+/+^, Tmem43^+/KI^* and *Tmem43^KI/KI^* (5 month). There was no difference among the genotypes. **c**, Co-IP data show that both TMEM43 WT and TMEM43-p.R372X are immuno-pulled down with Cx26 and Cx30. **d**,**f**,**h** Representative I-V curve measured from GLSs of control (black), carbenoxolone treated (**d**), GdCl_3_ treated (**f**) and pH6 treated (**h**) (red) cochlea. **e**,**g**,**i** Summary bar graph of **d** (**e**), **f** (**g**) and **h** (**i**). Glass pipette was filled with K-Gluconate internal solution. **j**, Representative I-V curve measured from naïve (black), control shRNA treated (green) and *Tmem43* shRNA treated (red) cochlea. **k,** Summary bar graph of (**j**). **l**, Representative current-voltage relationship of *Tmem43^+/+^* (red), *Tmem43^+/KI^* (green) and *Tmem43^KI/KI^* (blue). **m**, Summary bar graph of GLS currents from (**l**). **n**-**p**, GLS current measured from *Tmem43* KI shRNA un-infected and infected cochlea of *Tmem43^+/+^* (**n**), *Tmem43^+/KI^* (**o**) and *Tmem43^KI/KI^* (**p**) mice. Note that the *Tmem43* KI shRNA rescues impaired passive current of *Tmem43^+/KI^*. **q**, Summary bar graph of (**n-p**).

It has been reported that K^+^ channel expression is low in GLSs so most of their resting membrane conductance is mediated by gap junctions, and gap junction blockers or isolation of GLSs increase their membrane resistance dramatically^27^, implying that gap junction channels mediate passive conductance current in GLSs. To test a possible involvement of TMEM43 in connexin channel function, we performed whole-cell patch clamp in GLSs from acutely dissected cochlea tissue by whole-cell patch clamp. Indeed, the large passive current with a linear current-voltage (I-V) relationship was recorded from GLSs, which was abolished when gap junction channel blocker carbenoxolone was treated (**Fig. 6d,e**). As the GLS gap junction networks are known to take part in recirculation of cochlear K^+^ ions^1^, we treated a broad-spectrum non-selective cation channel blocker GdCl_3_^28^. Treatment of GdCl_3_ (30 μM) also abolished the passive current of GLSs (**Fig. 6f,g**). We next examined if pH change can gate the passive conductance of GLS as some connexin channels are reported to be pH sensitive^1^. When extracellular pH was lowered to 6, the passive conductance current was completely abolished (**Fig. 6h,i**). These results indicate that GLSs display the large passive conductance current which is cationic and pH sensitive. The similar elimination of the passive conductance current was observed when cultured cochlea tissue was infected with a virus carrying *Tmem43* shRNA (**Fig. 6j,k**). We subsequently measured the passive conductance current from *Tmem43^KI^* mouse. The passive conductance current from GLSs of *Tmem43^+/+^* was significantly reduced by 89% in *Tmem43^KI/KI^* mouse (**Fig. 6l,m**), indicating that TMEM43 contributes majorly to the passive conductance of the GLSs. The heterozygote mouse, *Tmem43^+/KI^*, which mimics the counterpart human ANSD subjects, showed a significant reduction of 63% in the passive conductance current in GLSs (**Fig. 6l,m**), confirming the significant pathogenic effect of the TMEM43 variant on the function of GLSs. To rescue the impaired TMEM43-p.R372X-mediated current, we generated a shRNA specifically targeting the KI sequence of *Tmem43^KI^* mouse. This *Tmem43* KI shRNA targets only KI allele but not WT allele (**Extended Data Fig. 4c,d, Fig. 6n,q**). When *Tmem43^+ KI^* heterozygote cochlea culture was infected with the virus carrying *Tmem43* KI shRNA, the reduced passive conductance current was fully rescued to the level comparable to *Tmem43^+/+^* (**Fig. 6o,q**) but not in *Tmem43^KI/KI^* homozygote cochlea culture (**Fig. 6p,q**). These results clearly explains the autosomal dominant effect of TMEM43-p.R372X and emphasize the role of TMEM43 as an essential component of connexin channels in mediating the passive conductance current in GLSs.

### Cochlear implant restores the impaired speech discrimination in the ANSD subjects

Clinically, determining the exact lesion site of hearing loss is critical for choosing a proper treatment. For example, subjects with conductive hearing loss or moderate sensorineural hearing loss due to limited damages to OHCs would benefit from hearing aids, whereas subjects with severe to profound sensorineural hearing loss, as a result of significant damages to OHCs or IHCs, would require a cochlea implant (CI). A subset of sensorineural hearing loss mainly affecting more central structures, such as spiral ganglion neuron or cochlear nerve, or central hearing loss originating from the brainstem or brain, may benefit from brainstem implant, but not from either hearing aids or CI. Because the main damage in *Tmem43^+KI^* mouse was restricted to GLSs, but not to the hair cells or spiral ganglion neurons at minimum 6~7 months of age, we could predict that subjects from SB162 could benefit from CI. Consequently, CI was performed on #284 and #304, even though the PTA of #284 did not meet a conventional criterion of CI (PTA equal to or exceeding 70dB^29^). Postoperatively, the open-set speech understanding of #284 with a deaf duration of 15 years more rapidly restored than that of control adult cochlear implantees (n=39), as measured by speech evaluation test tool^30,31^ (**Fig. 7a,b**). However, the restoration of speech discrimination ability in #304 with a longer deaf duration (about 25 years) was not as rapid as that in #284 (**Fig. 7c**). To interest, the responsiveness of cochlear nerve to stimuli of #304, measured by intracochlear electrically evoked ABR (e-ABR), was restored immediately after cochlear implantation **(Fig. 7d)**, which sharply contrasted with other ANSD subject with a postsynaptic etiology (**Fig. 7e**). These results suggest that limited improvement of speech discrimination in #304 is presumably due to cortical damage related with longer deaf duration, but not due to cochlear implant-related issues. Based on our results, early cochlear implantation may be recommended despite substantial residual hearing, in the case of TMEM43-p.R372X among adult onset, progressive ANSD subjects.

**Figure 7.**
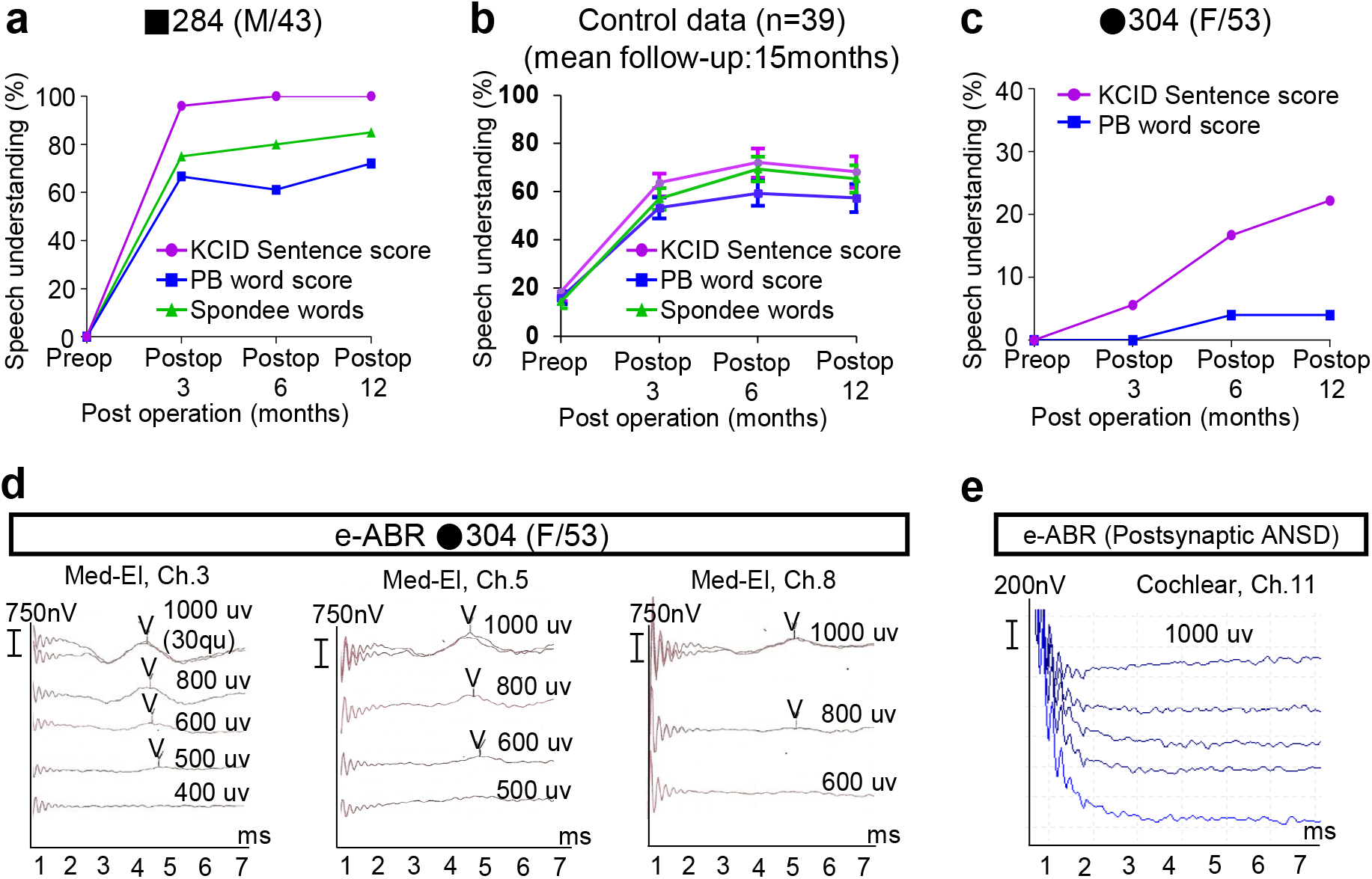
Timely cochlea implantations in family SB162 rescue impaired speech discrimination. **a**, Speech discrimination performance scores of subject 284 measured at 3, 6 and 12 months following cochlear implantation. Telephone speech discrimination was 100% at 12 months following cochlea implantation. **b**, Longitudinal change of postoperative auditory performance from control subjects (n=39) with adult onset progressive hearing loss: Scores at postoperative 3 months, 6 months, 1year as measured by K-CID, Spondee and PB word are shown. **c**, Subject 304 showed improved K-CID (up to 22.2%) and PB word (up to 4%) score after cochlear implantation. KCID Sentence score: Korean Central Institute for the Deaf score. PB word score: phonetically balanced word score. Spondee words: audiology a word with 2 syllables–disyllabic, that is pronounced with equal emphasis the 1^st^ and 2^nd^ syllables. **d,** The responses of Electrical Auditory Brainstem Responses (e-ABR) obtained through the cochlear implant electrodes inserted in the right ear (shown in red) are evident at the entire channel tested (No. 3, 5, 8) in subject 304. Conventionally, the responses of the right and left ear are shown in red and blue, respectively. The wave V (annotated on the waves) was detected at about 4ms after electrical stimulation and the thresholds was ranged from 500 to 800 μV. **e,** The results of e-ABR obtained through the electrodes implanted in the left ear of the representative case of postsynaptic auditory neuropathy spectrum disorder are shown in blue. The wave V was not detected even with the maximum intensity of stimulation (1000 μv) due to the dys-synchronic response of cochlear nerve.

## Discussion

In this study, we have identified a novel deafness locus, Chr 3: 13,165,401 −14,502,000 (3p25.1) and a new deafness gene, *TMEM43*, which, if altered, can cause ANSD in adults. The p.R372X variant of *TMEM43* is considered clearly pathogenic and causative of ANSD of HN66 and SB162, based on the guideline for the interpretation of classifying pathogenic variants (PS3, PM2, PP1_Strong and PS4_Supporting)^19,20^. Detection of this variant from two unrelated families (each from China and South Korea) sharing rare and characteristic ANSD phenotype significantly increases causality between this variant and ANSD. Unbiased exome sequencing screening was performed both for HN66 and SB162 in a parallel fashion with linkage analysis to minimize errors from spurious linkage. Significant overlap of linkage interval between HN66 and SB162 in Chr 3: 13,165,401 −14,502,000 made it least likely for our linkage data to be reflection of spurious linkage. Further, we have identified and characterized TMEM43 as an essential component of connexin channels contributing to the passive conductance current in GLSs which is necessary for hearing and speech discrimination. Until now, only the pathological functions of TMEM43 has been implicated in arrhythmia^22,23^ and tumor progression^32^. The variant p.S358L of *TMEM43* has been reported to cause ARVC by decreasing the expression and localization of tight junctions, redistributing the gap junction proteins from the surface to the cytoplasm, and decreasing the conduction velocity^23^.

Our study firstly demonstrates the presence of the CBX-, GdCl_3_- and low pH-sensitive passive conductance current in the GLSs of intact cochlea. The high expression of passive conductance channels, which gives rise to very low membrane resistance and linear I-V relationship, is known as the unique property that defines mature astrocytes in the brain^33^. Therefore, our findings imply that GLSs display similar properties as mature astrocytes. Consistently, GLSs express GFAP and GLAST^34^, which are common markers for mature astrocytes. In the brain, passive conductance channels are thought to be involved in K^+^ ion homeostasis^35^ and volume regulation^36,37^. In the cochlea, the endocochlear potential of +80mV is generated by maintaining high K^+^ concentration in the endolymph, and this potential is critical for activation of hair cells^38^. The presence of passive conductance current implicates an important role of GLSs in homeostasis of K^+^ and volume regulation. We have observed a significantly smaller inner border cell size of *Tmem43^KI^* mice compared to control, with loss of passive conductance current. We hypothesize that the smaller inner border cells have smaller capacity for K^+^ uptake. The loss of the passive conductance current indicates impaired recycling of K^+^. Such breakdown of K^+^ homeostasis would make hair cells difficult to depolarize, resulting in a disruption of speech discrimination ability.

It is of interest that the onset time of pathology and hearing loss in heterozygous *Tmem43^KI^* mouse (6~7 months of age which equals mid-thirties in human) precisely matches the onset time of a full extent of ANSD (mid-thirties) in our affected subjects. This raises a fundamental question of why canonical ANSD does not occur in young age. The passive conductance current recordings from the cochlea of P5-7 pups revealed a 63% reduction in passive conductance current in heterozygous *Tmem43^KI^* mice. We also showed that lowering the pH to 6 reduced the passive current of GLS. The late-onset hearing loss could be explained by a reduction of cochlear pH in aging mouse and human, similar to what happens in aging brain^39^. The normal hearing in heterozygote *Tmem43^KI^* mouse and the ANSD subjects at young age could be owing to the remaining passive conductance current that might be enough to maintain homeostatic functions of GLSs. In contrast, the hearing loss in the aging *Tmem43^KI^* mice and ANSD subjects can be explained with further reduction of the remaining current due to the low pH-sensitivity, impairing the K^+^ homeostatic in GLSs. These exciting possibilities await future investigations.

Our study provides a potential new prognostic genetic marker *TMEM43-*p.R372X for ANSD patients. More importantly, our study suggests that if a causative gene of ANSD is expressed mainly at GLSs, it could also be used as a favorable prognostic marker for CI. Although CI successfully rehabilitates severe to profound sensorineural hearing loss in many cases, especially adult ANSD patients usually hesitate to receive the surgery due to uncertainty of surgery outcome regarding speech discrimination, side-effects including loss of residual hearing, and high cost. Given this, an ANSD-causative variant of GLS-specific gene such as p.R372X of *TMEM43* could serve as a potential indicator of positive CI surgery outcome if the surgery is timely done. In this sense, our study also proposes the optimal time of CI surgery: #284 who received CI at the age 43 recovered his sentence understanding ability to 100% (**Fig. 7a**), while #304 who received CI at the age 53 recovered her ability to 22.2% one year after surgery (**Fig. 7c,d**). Relatively slow functional recovery from #304 could be attributed to the cortical damage due to longer deaf duration. Thus, we may have to recommend the ANSD patients carrying *TMEM43-p.R372X* to receive a CI surgery as soon as they start to experience significantly diminished speech discrimination even though they retain significant residual sound detection ability. Our study provides an important step towards identifying more *TMEM43*-related ANSD patients, enabling further study with larger number of subjects carrying this allele.

In conclusion, we have characterized TMEM43 as an essential component of connexin channels and have delineated that an alteration of this protein causes functional and morphological abnormalities in GLSs, resulting in the failure to maintain speech discrimination in aged human. Through these mechanistic insights, we have elucidated a link between abnormality in cochlear GLSs and impaired human speech discrimination. We further provide a model platform in which the personalized timing and mode of auditory rehabilitation can be determined, highlighting the importance of a precision medicine-based approach.

## Methods

### Family data

Clinical examination was performed on a total of 18 subjects: 6 male and 12 female subjects whose age ranged between 22 and 84 years from family SB162. Five affected subjects with progressive “ANSD” and 11 unaffected subjects in a 5 generation Chinese Han HN66 family participated in the present study. Informed consent was obtained from all participants in this study. Pure-tone audiometry was carried out to evaluate the air and bone conduction thresholds. The binaural mean pure-tone average of thresholds for air conduction (in dB SPL) was calculated for audiometric frequencies of 0.25, 1, 2, and 4 kHz (PTA 0.25, 1, 2, 4kHz). The speech discrimination score was measured whenever possible. Hearing impairment was additionally assessed by the means of ABRs and DPOAEs. The ABR threshold is a minimal amplitude of sound that can elicit the wave V response. Otoscopic examination and tympanometry with acoustic reflex testing were performed systematically to rule out any conductive hearing impairment. To evaluate the function of outer hair cells, sound pressure of DPOAE and surrounding noise were evaluated using a SmartEP (IHS, Miami, FL). An Etymotic 10B+ microphone was inserted to measure sound pressure in the external ear canal, and pure tone from 3.6 to 14.4 kHz was presented using Etymotic ER2 stimulator. Frequencies were acquired with an F2-F1 ratio of 1.22. Stimulation levels ranged from 65 to 25 dB SPL at 10-dB interval. This study was approved by the institutional review board of Seoul National University Bundang Hospital (IRB-B-1007-105-402) and the ethics committee of Xiangya Hospital of Central South University (ID: 201603518).

### Linkage analysis

The Infinium Global Screening Array panel (Illumina, San Diego, CA), consisting of 660 thousands of SNPs, was used to conduct a whole genome linkage scan with 18 subjects (10 affected and 8 unaffected) in SB182 and 9 subjects (5 affected and 4 unaffected) in HN66. The genomic DNAs were extracted from the peripheral blood. Multipoint log_10_ *Odds* (LOD) scores were computed using MERLIN (version 1.1.2) and was given for all genotyped positions. Analyses were performed assuming the dominant traits based on family history. The disease allele frequency was set to 0.0001, and the penetrance for homozygous normal, heterozygous, and homozygous affected were set to 0.0001, 1.0, and 1.0 respectively. Family members with unclear status were coded as unknown. Because Merlin allows a maximum of 24 bits for a family by default, the family was split into sub-families having common ancestors in SB162.

### Targeted exome sequencing and exome sequencing

Targeted exome sequencing of the critical region 3p25.1 from the linkage analysis for four subjects (SB162-#284, #289, #290, #291) was performed and the candidate variants were listed (**Table 1**). The variants < 1% minor allele frequency in ESP6500 and 1000G were initially identified. Only inheritance pattern-matched variants were remained, and the variants in dbSNP, but not in flagged dbSNP were filtered out. We used our in-house database having KRGDB_1722 individuals to remove Korean-specific common variants. In addition to single nucleotide variation (SNV), possible presence of copy number variations (CNV) was also identified using Excavator^8^. A genome level log2 ratio for chromosome 3 was investigated. The subjects proband was screened for hotspot variants causing Hearing Loss in Korean and Chinese population via hot spot screening, which include *GJB2, SLC26A4* and mitochondrial 12S rRNA. In parallel, being blind to the linkage analysis results, we also performed exome sequencing on the four subjects (SB162-#284, #289, #290, #291) in SB162 and 2 affected subjects (#36 and #28) and 1 unaffected subject (#22). Bioinformatics analyses were performed as previously described^8^. Thereafter, the final candidate variants in these families were verified using Sanger sequencing. The variants in non-coding regions as well as synonymous variants in coding regions were filtered out. Variants with a minor allele frequency of less than 1% were selected based on the Exome Sequencing Project 6500 (ESP6500), 1000 Genome Project (1000G), Exome Aggregation Consortium (ExAC), and our in-house database containing the exomes of 81 Korean individuals. Based on the autosomal dominant inheritance pattern, single heterozygous variants with sufficient read depths (>10X) and genotype quality (>20) were selected when they were commonly found in all the affected siblings. Further, variants of dbSNP ID with no clinical significance were filtered out. Finally, 20 possible candidate variants in SB162 and 5 candidates in HN66 that co-segregated with ANSD phenotype remained (**Table 1** and **Supplementary Table 1**). To ascertain the candidate pathogenic variants, ES was performed on two affected subjects (#36 and #28) and one unaffected subject (#22) and the resulting sequences were processed using the pipeline that we have previously described^40^. After initial quality control, variants were firstly filtered using the deafness-associated gene list (DAGL) to search for variants in known deafness genes. Then, the remaining variants were filtered according to the criteria as follows: (1) locating at the coding exons and intron-exon adjacent regions; (2) MAF < 0.1% in multiple databases including the Exome Aggregation Consortium (EXaC) (http://exac.broadinstitute.org/), the 1000 Genome Project (http://browser.1000genomes.org), gnomAD (http://gnomad.broadinstitute.org/) and the ATCG (Annoroad Typical Chinese Genomes) database; (3) heterozygous variants in the form of non-synonymous, non-sense, splice sites, Indels, codon mutations; (4) predicted pathogenicity for missense mutations using SIFT (http://sift.jcvi.org), MutationTaster (http://mutationassessor.org), and Polyphen2 (http://genetics.bwh.harvard.edu/pph2). Candidate pathogenic variants were further conformed by mapping the filtered variants obtained from WES to the linkage regions.

### Whole Genome Sequencing

Whole genome sequencing was performed on 8 DNA samples (SB162-284, 289, 290, 304, 307, 309, 324, 332) using HiSeq2000. Bioinformatics analysis (alignment to the hg19 reference genome, deduplication, local de novo assembly, and variant calling) was performed using BWA, picard, and GATK). Variants firstly were restricted to those in specified regions (chr3:11,883,000-chr3:14,502,000, based on the result of linkage analysis). We then filtered out the variants that were not matched to inheritance patterns, and that were in Korean reference database (KRGDB, KOVA and in-house Samsung Genome Institute control subjects database) and the 1000G project.

### Evaluation of speech discrimination

Speech performance was evaluated preoperatively, and 3, 6, and 12 months after CI using Korean version of the Central Institute for the Deaf (K-CID) and the Korean PB word by experienced speech-therapist. The Korean PB word included 40 monosyllabic words that are phonetically balanced, and K-CID was composed of sentences to evaluate speech discrimination. K-CID and PB word were presented in sound-field with condition of audio-only and no visual cues. Patients were instructed to repeat the speech stimulus verbally, and the score was measured with the correction ratio (%).

### Animals and housing

*Tmem43*-R372X Knock-in (C57BL/6J;129S-*Tmem43^tm1Cby^*) mice, 129Sv/Ev mice and crab-eating macaque were used. All animals were kept on a 12 hours light-dark cycle in a specific-pathogen-free facility with controlled temperature and humidity and had free access to food and water. All experimental procedures were conducted according to protocols approved by the directives of the Institutional Animal Care and Use Committee of Seoul National University Bundang Hospital (Seongnam, Republic of Korea) and the Institutional Animal Care and Use Committee of IBS (Daejeon, Republic of Korea). As the sequence difference between *Tmem43^+/+^* and *Tmem43^KI/KI^* was too small to be detected by conventional PCR method, mouse cDNA was amplified with primers (F: 5-gtttatgggcctcaacctcatg-3, R: 5-caggcttcactccagctttttgg-3) and then enzyme digested with BsiEI (Thermofisher #FD0894). Genotyping result using this method is shown in Fig. 5a. The expected band size for WT is 90bp and123bp, and for KI/KI is 213bp.

### Construction of *Tmem43-R372X* knock-in mice model

The *Tmem43*-R372X Knock-in (C57BL/6J;129S-*Tmem43^tm1Cby^*) mouse model was generated using Clustered Regularly Interspaced Short Palindromic Repeats (CRISPR) technology. In such a model, Arginine (R) was replaced with the STOP codon at position 372 of the TMEM43 protein (gene ID: NM_028766). The criteria used for gRNA selection are the distance to the modification site and the off-target profile. Based on these guidelines, two gRNA candidates (*Tmem43* gRNA1:5’-TTAGAGCAGCCCACAGCGGTCGG-3’; *Tmem43* gRNA2: 5’-CCGATTAGAGCAGCCCACAGCGG-3’), located in proximity to the mutation site, were selected and evaluated. To determine gRNA activity, a SURVEYOR assay was performed, using SURVEYOR mutation detection kit (IDT), according to the manufacturer’s instructions. Based on the validated gRNAs, a single stranded oligodeoxynucleotide donor (ssODN) was designed and synthesized. A 160 bp donor contains two homology arms flanking introduced (C ➔ T) mutation in exon 12 corresponding to 1114 position in the *Tmem43* gene. Moreover, to protect the repaired genome from being re-targeted by Cas9 complex, additional mutations were introduced corresponding to a second stop codon. The 62 C57BL/6J embryos were injected through a cytoplasmic route with a CRISPR cocktail containing ssODN donor, gRNA transcripts *Tmem43* and Cas9 mRNA. Forty-nine out of 62 embryos passed the quality screening and were implanted into two surrogate CD1 mice. From this round of microinjection, all females became pregnant, and gave birth to nine pups F0. To confirm the germline transmission of the transgene, female founder F0 was mated with the wild type C57BL/6J male and subjected to F1 breeding to confirm germline transmission of the transgene. All animals were analyzed by PCR and sequencing for the presence of correct point mutation at the target site. For genotyping, genomic DNA was extracted from 1-to 2-mm-long tail tips using the DNeasy^®^ Blood & Tissue kit (Qiagen, Hilden, Germany). Genomic DNA (5 *μ*l) was analyzed by PCR in a final volume of 50 *μl* in the presence of 25 mM MgCl_2_ and 2 *m*M dNTPs, at 10 pM of each primer, and 0.02 U of TOYOBO KOD Hot Start DNA polymerase (Invitrogen/Life Technologies, Billerica, Massachusetts, USA) with primers: Forward (5’-CCACAGTGGACTGGTTTCCT-3’) and Reverse (5’-GGCTTCACTCCAGCTTTTTG-3’) detecting the presence of the knock-in allele (213 base pairs). After a denaturing step at 95°C for 2 minutes, 35 cycles of PCR were performed, each consisting of a denaturing step at 95°C for 20 seconds, followed by an annealing phase at 59°C for 10 seconds and an elongation step at 72°C for 10 seconds. PCR was finished by a 10-minute extension step at 72°C. Amplified products were confirmed by Sanger sequencing (Macrogen Inc., Seoul, KOR).

### Plasmids

TMEM43 (Myc-DDK-tagged)-Human transmembrane protein 43 (TMEM43) (GenBank accession no. NM_024334.2) was purchased from OriGene (RC200998) and cloned into CMV-MCS-IRES2-EGFP vector using BglII/XmaI sites. hTMEM43-p.R372X mutation and p.401X mutation (to put stop codon before FLAG) were obtained by performing oligonucleotide-directed mutagenesis using the EZchange site-directed mutagenesis kit (EZ004S, Enzynomics). mTMEM43 (GenBank accession no. NM_028766.2) was obtained by using a RT–PCR based gateway cloning method (Invitrogen) and cloned into CMV-EGFP-C1 vector using BglII/SalI sites. GJB2 (NM_004004) Human Tagged ORF Clone was purchased from OriGene (RC202092) and Cx30-msfGFP was purchased from Addgene (69019).

### Heterologous expression of TMEM43 on HEK293T cell lines

Human embryonic kidney (HEK) 293T cells were purchased from ATCC (CRL-3216). The cell line has been tested for mycoplasma contamination. HEK293T cell was cultured in DMEM (10-013, Corning) supplemented with 10% heat-inactivated fetal bovine serum (10082-147, Gibco) and 10,000 units/ml penicillin–streptomycin (15140-122, Gibco) at 37°C in a humidified atmosphere of 95% air and 5% CO_2_. Transfection of expression vectors was performed with Effectene Transfection Reagent (Effectene, 301425, Qiagen), according to the manufacturer’s protocol. One day prior to performing the experiments, HEK293T cells were transfected with each DNA 1.5μg per 60mm dish. Ratio of DNA to Effectene Reagent is 1:10.

### RT-PCR

The 129Sv/Ev mice were deeply anesthetized with pentobarbital sodium (50 mg/kg). Total RNA was extracted from their whole cochlea (P1), heart (P120), eye (P42), brain (P28), kidney (P28), and liver (P28) to analyze the expression of the target mRNAs using TRIZOL^®^ (Gibco BRL, Gaithersburg, MD, USA) and a column from an RNeasy Mini Kit (Qiagen, Valencia, USA), performed in accordance with the manufacturer’s instructions. The RNA obtained was used to synthesize cDNA with oligo (dT) primers, one-twentieth of which was used for one PCR reaction. The DNA amplification was performed in a final volume of 30 μl. PCR cycling conditions were 95°C for 2 min, followed by 35 cycles of 95°C for 20 s, 55°C for 10 s, and 70°C for 10 s, with a final step of 72°C for 5 min. Amplified product (15 μl) was separated using electrophoresis in a 2% agarose gel and visualized with ethidium bromide. We analyzed the transcripts of TMEM43 and GAPDH by primer pairs, as follows: the forward primer (5-CTTCCTGGAACGGCTGAG-3’) and the reverse primer (5’-CACCAGCCTTCCTTCATTCT-3’) for *Tmem43*, and the forward primer (5’-ACCACAGTCCATGCCATCAC-3’) and the reverse primer (5’-CACCACCCTGTTGCTGTAGCC-3’) for *Gapdh*.

### Immunocytofluorescence on cochlear tissue

Primary antibodies used are rabbit anti-TMEM43 monoclonal (1:500, Ab184164, Abcam) (Ab_1-1_), rabbit anti-TMEM43 polyclonal (1:100, NBP1-84132, Novus) (Ab_1-2_), mouse anti-mCherry monoclonal (1:500, ab125096, Abcam), mouse anti-calretinin monoclonal (1:500, MAB1568, Millipore), goat anti-Na^+^, K^+^-ATPase-α3 polyclonal (NKA, 1:250, SC-16052, Santa Cruz), and mouse anti-CtBP2 monoclonal (1:500, 612044, BD Transduction Laboratories) antibodies. Ab_1-1_ was used in cultured cochlea and Ab_1-2_ was used in freshly dissected cochlea. Secondary antibodies generated either in donkeys or goats (life technologies) were used at 1:1000. Secondary antibodies used are donkey anti-mouse alexa fluor 555 (A-31570, Thermo fischer scientific), donkey anti-rabbit alexa fluor 488 (A-21206, Thermo fischer scientific), donkey anti-goat alexa fluor 633 (A-21082, Thermo fischer scientific), goat anti-rabbit alexa fluor 555 (A-21428, Thermo fischer scientific), goat anti-mouse alexa fluor 488 (A-11001, Thermo fischer scientific), alexa fluor 488 goat anti-rabbit IgG (H+L) (A11008, Life technologies). No immunoreactivity was found when the primary antibodies were omitted. Mouse inner ears (C57BL/6) at various time points postnatally and the cochlea from three monkeys weighing 5kg were fixed in ice cold 4% paraformaldehyde, for 1 h. Cochlear turns were carefully excised and incubated in blocking/permeabilizing buffer (PBS with goat or donkey serum and 0.25% Triton X-100). Then, the preparations were incubated overnight at 4°C with primary antibodies diluted in the blocking/permeabilizing buffer. After 3 washes, the cochlear turns were reacted with fluorescence labeled secondary antibodies diluted in blocking/permeabilizing buffer for 1 h at room temperature. Samples were then rinsed once with blocking/permeabilizing buffer and twice with PBS. Using Fluorsave reagent (Calbiochem, 345789), the tissues were mounted on glass slides and covered with coverslip. Specific immunolabeling was initially examined under epifluorescence microscope and high-resolution images were obtained using a confocal laser scanning microscope (LSM710, Zeiss).

### Immunocytofluorescence on heterologous system

For immunocytochemistry, hTMEM43-IRES2-EGFP was transfected into HEK293T cells one day prior to staining. Cells were fixed in 4% paraformaldehyde for 30 min at room temperature and washed 3 times with PBS. The permeabilized group contained 3% Triton X-100 in the blocking solution with 2% goat serum and 2% donkey serum but the impermeabilized group excluded Triton X-100 in the blocking solution. Cells were incubated with rabbit anti-TMEM43 polyclonal antibody (1:100, NBP1-84132, Novus) (Ab_1-2_) and mouse anti-FLAG (1:500, F1804, Sigma) for overnight. After washing, goat anti-Rabbit IgG Alexa Flour 647 (1:1000, A21429, Invitrogen) and donkey anti-Mouse Alexa 555 (1:1000, A31570, Molecular Probe) were added and incubated for 2 hours at room temperature. The cells were washed 3 times and mounted, and then observed under a Nikon A1 confocal microscope.

### In situ hybridization

To make specific riboprobe for mRNA of Tmem43 and Spmx, we cloned partial cDNA fragments of Tmem43 and Spmx. Primers were as follows: Tmem43, forward: 5’-TGTTCGTGGGGCTAATGACC-3; reverse: 5’-TAGTTGGTCACCTCGTTGCC-3’, T7-reverse primer: 5 ‘-TAATACGACTCACTATAGGGAGATCACAGTAACCACGTGAGCC-3’; Spmx, forward: 5 ‘-TTTCCACACGGTCAAGCCTTT-3; reverse: 5’-TCTCCTCAAAACCACACTTCCC-3’, T7-reverse primer: 5’-TAATACGACTCACTATAGGGAGAGTCTTGGGCATTCAGGAGGTT-3’. The plasmid was linearized and used for in vitro transcription (Roche Dignostics, Indianapolis, IN, USA) to label RNA probes with digoxigenin-UTP. The 3’ and 5’ – DIG labeled miRCURY LNA miRNA Detection probes (Qiagen, negative control Scramble-miR cat. # YD00699004-BCG) were used for In situ hybridization. In situ hybridization was performed as previously described with some modifications. Frozen cochlea was sectioned at 20 μm thickness on a cryostat. The sections were then fixed in 4 % paraformaldehyde, washed with PBS, and acetylated for 10 min. The sections were incubated with the hybridization buffer (50 % formamide, 4X SSC, 0.1 % CHAPS, 5 mM EDTA, 0.1 % Tween-20, 1.25 × Denhartdt’s, 125 μg/ml yeast tRNA, 50 μg/ml Heparin) and digoxigenin-labeled probes (200 ng) for 18 h at 60 °C. Non-specific hybridization was removed by washing in 2X SSC for 10 min and in 0.1X SSC at 50 °C for 15 min. For immunological detection of digoxigenin-labeled hybrids, the sections were incubated with anti-digoxigenin antibody conjugated with alkaline phosphatase (Roche Diagnostics) for 1 h, and the colour reaction was carried out with NBT/BCIP (4-nitroblue tetrazolium chloride/bromo-4-chloro-3-indolyl phosphate; Sigma). Sections were dehydrated and mounted with Vectamount (Vector Laboratories, Burlingame, CA, USA).

### Cochlear organotypic culture and shRNA treatment

Neonatal mouse (C57/BL6, P4) cochlear turns were isolated in ice cold sterile Hanks’ Balanced Salt Solution (HBSS). Cochlear segments were attached on Cell-Tak (354240, Corning) coated coverslips and incubated overnight in DMEM/F12 medium containing 1% fetal bovine serum, 5μg/ml ampicillin, B27 and N2 (37 °C, 5% CO_2_ humidified incubator). Upon confirming stable attachment, the tissues were treated with either scramble or shRNA diluted in culture medium (1:1000) for 48 h. The medium was then replaced with a fresh one. After additional 48 h of incubation, the tissues were subjected to further examination.

### Construction and packaging of shRNA–lentiviral vectors

For TMEM43 gene silencing, candidate for both mouse and rat *Tmem43* shRNA (GenBank accession no. NM_028766.2 and NM_001007745.1) sequences were cloned into lentiviral pSicoR vector using XhoI/XbaI sites, as previously described^41^. Knockdown efficiency was determined by a reduction in fluorescence expression of CMV-EGFP-TMEM43 when overexpressed with *Tmem43* shRNA candidate containing pSicoR-*Tmem43* shRNA-mCh vector. Validated shRNA candidate containing pSicoR vector was packaged into high-titer lentivirus by KIST virus facility (Republic of Korea). In brief, the lentiviral vectors were produced by co-transfecting each pSicoR vector with the ViraPower lentiviral packaging mix (Invitrogen) in the 293FT packaging cell line. The supernatant was collected and concentrated by ultracentrifugation. For efficient shRNA delivery, lentiviral vectors were always produced at subneutral pH (≤pH 7.0). The target regions of *Tmem43* shRNA is 5’-GCTCTTGTCTGACCCAAATTA-3’ and *Tmem43* KI shRNA is 5’-CTCTTCTACTGATGACTGTGG-3’. For control shRNA, scrambled sequence 5’-TCGCATAGCGTATGCCGTT-3’ was inserted in place of shRNA sequence.

### Chemicals

Chemicals used in this study were all purchased from Sigma-Aldrich.

### Electrophysiological recording in cochlear supporting cells

Cochlea of P5-P7 *Tmem43-* p.R372X Knock-in mouse pups were isolated in HEPES buffer containing (mM): 144 NaCl, 5.8 KCl, 1.3 CaCl_2_, 2 MgCl_2_, 10 HEPES, 0.7 NaH_2_PO_4_ and 5.6 D-glucose (pH 7.4 was adjusted with NaOH). All recordings were done with same HEPES buffer as external solution. Stria vascularis and tectorial membrane were carefully peeled off and the remaining cellular organization of the organ of Corti was left intact. Acutely dissected cochlea turn was used within 2 hours of dissection. Glia-like supporting cells that are located below inner hair cell layer were whole cell patch clamped and current traces were elicited by 1 sec ramps ascending from −100mV to +100mV with −60mV holding potential. Recording electrodes (7-11M Ω) supplemented with (mM): 126 KOH, 126 Gluconate, 5 HEPES, 0.5 MgCl_2_ and 10 BAPTA (pH adjusted to 7.3 with KOH) advanced through the tissue under positive pressure. Slice chamber was mounted on the stage of an upright Hamamatsu digital camera viewed with an X60 water immersion objective with infrared differential interference contrast optics using Imaging Workbench Software. Electrical signals were digitized and sampled with Digidata 1320A and Multiclamp 700B amplifier (Molecular Devices) using pCLAMP 10.2 software. Data were sampled at 10 kHz and filtered at 1 kHz. Glass pipette were pulled from micropipette puller (P-97, Sutter Instrument) and all experiments were conducted at a room temperature of 20-22°C.

### Cell surface biotinylation, co-immunoprecipitation and western blot

For biotinylation, CMV-hTMEM43 WT-IRES2-EGFP or CMV-hTMEM43 R372X-IRES2-tdTomato vector were transfected into HEK293T cells 1 day prior to the experiment day. Transfected cells were washed three times with PBS and cell surface-expressed proteins were biotinylated in PBS containing Ez-link sulfo-NHS-LC-Biotin (21335, Thermo) for 30 min. After biotinylation, cells were washed with quenching buffer (100mM glycine in PBS) to remove excess biotin and then washed three times with PBS. The cells were then lysed and incubated with high capacity NeutrAvidin-Agarose Resin (29204, Thermo). After three washes with lysis buffer, bound proteins are eluted by the SDS sample buffer and subjected to western blot analysis. Primary antibody used is: rabbit anti-TMEM43 polyclonal (1:100, NBP1-84132, Novus) (Ab_1-2_). Secondary antibody used is: Donkey anti-rabbit HRP (NA9340, Amersham). For CHX assay, 40 ug/ml CHX was treated to TMEM43 WT and p.R372X expressing HEK293T cells and analyzed in 3 hours interval. Anti-FLAG M2 (1:2000, F1804, Sigma-Aldrich) was used for western blot. For co-immunoprecipitation, cell lysates were prepared in a buffer containing 50 mM Tris-HCl (pH 7.5), 150 mM NaCl, 1% NP-40, 10 mM NaF, and protease and phosphatase inhibitor cocktail. Equal amounts of precleared cell lysates were incubated with rabbit anti-FLAG (2368s, Cell Signaling) overnight. Protein A/G-Agarose beads (Thermo Fisher) were added to the mixtures and further incubated for 2 h, followed by a wash with lysis buffer. Bound proteins were eluted from the beads with SDS-PAGE sample buffer and western blotting was performed with rabbit anti-TMEM43 (Ab_1-2_).

### Scanning electron microscope (SEM) analysis

For SEM analysis, the organ of Corti were performed using a method previously described^42^. Briefly, the cochleae were immediately isolated from the euthanized *Tmem43^+/+^* and *Tmem43^+/KI^* mice at 2, 7, and 13 months of age and fixed in a solution of 2% PFA dissolved in 0.1 M sodium cacodylate buffer (pH 7.4) containing 2.5% glutaraldehyde for 1 hour at room temperature. The bony capsule was dissected out and the lateral wall, Reissner’s membrane, and tectorial membrane were removed. The organ of Corti was then dissected and fixed in a solution of 0.1 M sodium cacodylate buffer (pH 7.4), 2mM calcium chloride, 2.5% glutaraldehyde, and 3.5% sucrose for overnight at 4°C. Following fixation, samples were prepared for SEM using the osmium tetroxide – thiocarbohydrazide (OTOTO) method^42^. Then, the specimens were dehydrated in a graded ethanol solution, dried using a critical point dryer (HCP-2, Hitachi, Japan), attached on the stub, and then coated with platinum using a sputter coater (E1030, Hitachi, Japan). The surfaces of the organ of Corti were captured under a cold-field emission SEM (SU8220, Hitachi, Japan) that was operated at 10 or 15 kV. Micrograph measurements were performed using ImageJ software (National Institutes of Health, http://rsbweb.nih.gov/ij/), via the polygon tool for cell area measurements.

### Bioinformatics

In silico prediction Algorithm: Polyphen-2 (http://genetics.bwh.harvard.edu/pph2/index.shtml); SIFT (http://sift.jcvi.org/www/SIFT_chr_coords_submit.html); Conservation tools: GERP++ score in the UCSC Genome Browser (http://genome-asia.ucsc.edu/); ExAC, Exome Aggregation Consortium (http://exac.broadinstitute.org/); 1000 Genomes (https://www.ncbi.nlm.nih.gov/variation/tools/1000genomes/); KRGDB, Korean Reference Genome DB (http://152.99.75.168/KRGDB/); TOPMED, Trans-Omics for Precision Medicine (https://www.nhlbiwgs.org/). Secondary structure prediction of TMEM43 was done with ‘TMHMM v2.0’ server (http://www.cbs.dtu.dk/services/TMHMM-2.0/). TMEM43 multiple amino acid sequence alignment was done by ‘Clustal Omega version 1.2.4’ server (https://www.ebi.ac.uk/Tools/msa/clustalo/) and phylogenetic analyses were conducted in ‘MEGA version 5’^43^.

## Data analysis and statistical analysis

Off-line analysis was carried out using Clampfit version 10.4.1.10 and GraphPad Prism version 7.02 software. When comparing between two samples, significance of data was assessed by Student’s two-tailed unpaired t-test when samples showed normal distribution and assessed by Mann-Whitney test when samples did not pass normality test. Samples that passed normality test but not equal variance test were assessed with Welch’s correction. Comparing more than 2 samples were analyzed using one way ANOVA with Tukey’s post-hoc test when data passed normality test and Kruskal-Wallis test with Dunn’s post-hoc test when data did not pass normality test. Significance levels were given as: N.S. *P*>0.05, \**P*<0.05, \*\**P*<0.01, \*\*\**P*<0.001 and #*P*<0.0001.

## Acknowledgments

Authors thank Dr. Young-Woo Seo at Korea Basic Science Institute Gwang-Ju Center for his assistance in image analysis with Imaris software. This research was supported by a grant of the Korea Health Technology R&D Project through the Korea Health Industry Development Institute (KHIDI), funded by the Ministry of Health & Welfare, Republic of Korea (grant number: HI15C1632, and HI17C0952 to B.Y. Choi), Creative Research Initiative Program, Korean National Research Foundation (2015R1A3A2066619 to C.J. Lee) and the National Research Council of Science & Technology (NST) grant by the Korea government (No. CRC-15-04-KIST to S.J. Oh).

## Author contributions

H.R.P., J.W.K., S.H.O. and B.Y.C. recruited the family and provided samples. M.W.S. and X.L. did linkage analysis. N.K., J.L. and W.P. contributed to NGS. A.R.K., H.C. and X.L. did Sanger sequencing & NGS results analysis. J.H.H., S.S., X.C. performed Sanger sequencing from additionally recruited family members. M.Y.K., S.B. and Z-Y.C. maintained cell line and IHC. M.W.L. and H.W.S. performed RT-PCR and initial experiments on trafficking of our target protein in cell lines. J.B.S. made videos for localization of our target protein and also performed IHC of the mouse cochlea. J.J.H., B.J.K., D.H.W., E.Y. and K.S. performed dissection and IHC of the monkey cochlea. E.Y. and K.S. also contributed to making mouse cochlea culture, infecting it with TMEM43 shRNA and performing electrophysiology. K.Y.L., M.A.K., U.K.K. did SEM experiments. M.W.J. contributed to electro-physiology experiments, developed shRNA to knockdown *Tmem43* and did immunocytochemistry for TMEM43 expressed cells. S.J.O. L.M., C.H. contributed to cell surface biotinylation and western blot data. S.L and M.Y.K performed audiological testing of the mice. T.Y.K., A.N. and Z.H. performed coimmunoprecipitation and cycloheximide chase assay. D.Y.O. contributed to recollection and organization of the data into figures. J.J.H. also performed audiograms and speech evaluation, and B.Y.C., Y.F. and C. J. L planned experiments, wrote the manuscript and supervised the project.

