## Extended Figures for "Mutations in TMEM43 cause autosomal dominant auditory neuropathy spectrum disorder via interaction with connexin-mediated passive conductance channels"

#### Extended Fig.1

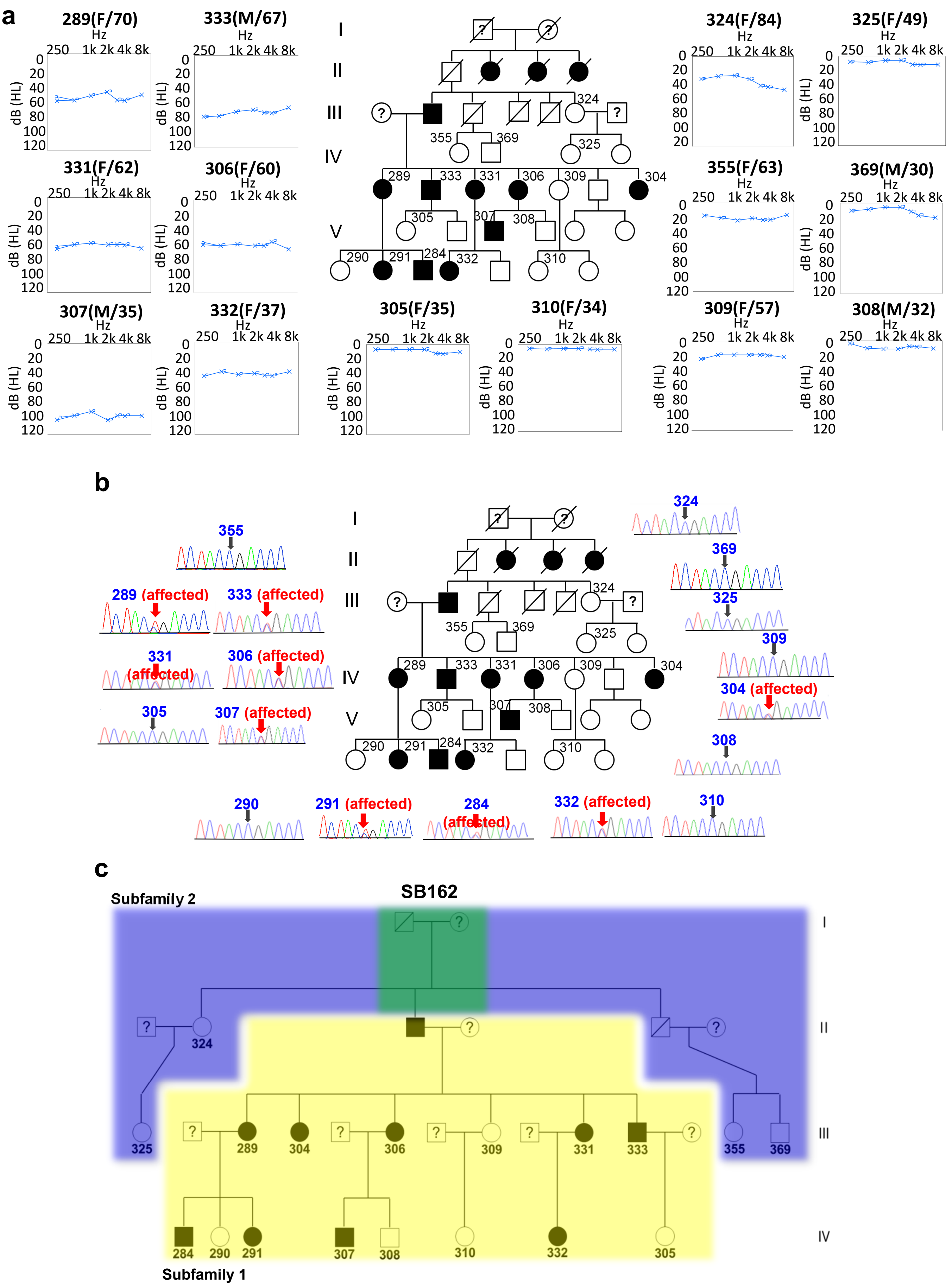

**Extended Data Fig. 1: A point mutation, p.R372X of TMEM43 is the cause of hearing loss in family SB162 as documented by linkage analysis, aCGH and segregation study among family members.**

**a,** Left six subjects (#289, #306, #307, #331, #332 and #333) show ANSD with 40dB or more degree of pure tone averages as calculated by averages of pure tone thresholds of 500Hz, 1kHz, 2kHz and 4kHz. At thirties, affected subjects lose hearing. The auditory thresholds of other subjects stay within a normal range. **c,** Subfamily 1 and 2 shaded in yellow and violet, respectively. A common ancestor shared by two subfamilies is shaded in green **b,** The trace from single heterozygous TMEM43 variant allele (p.R372X) is indicated in red on the chromatograms. Open and filled symbols indicate unaffected and affected subjects, respectively. Subject numbers are superscripted

#### Extended Fig.2

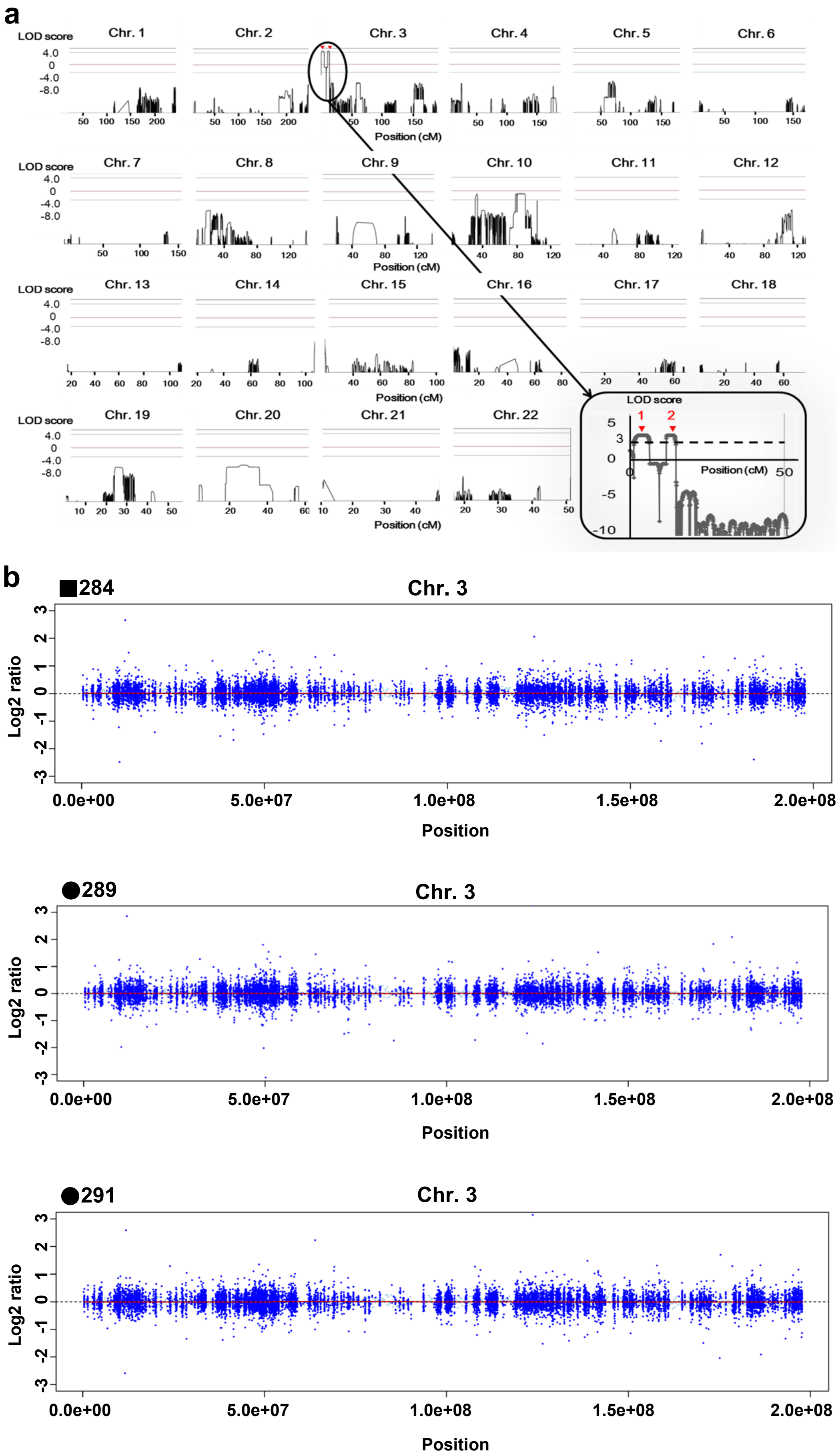

**Extended Data Fig. 2: TMEM43-p.R372X is mapped to chromosome3, without changing copy number variation.**

**a**, Merlin multipoint linkage analysis demonstrates LOD scores greater than 3.0 only on chromosome 3p25-26. Inset: magnified view of two regions on Chr. 3, spanning 6.7cM and 3cM, respectively. Red arrowhead 1: region #1 (Chr. 3: 1,946,000 - 5,956,000), Red arrowhead 2: region #2 (Chr. 3:11,883,000 - 14,502,000).
**b**, Presence of copy number variation checked by Excavator. Each panel shows a genome level log 2 ratio for chromosome 3. There is no significant copy number variation.

#### Extended Fig.3

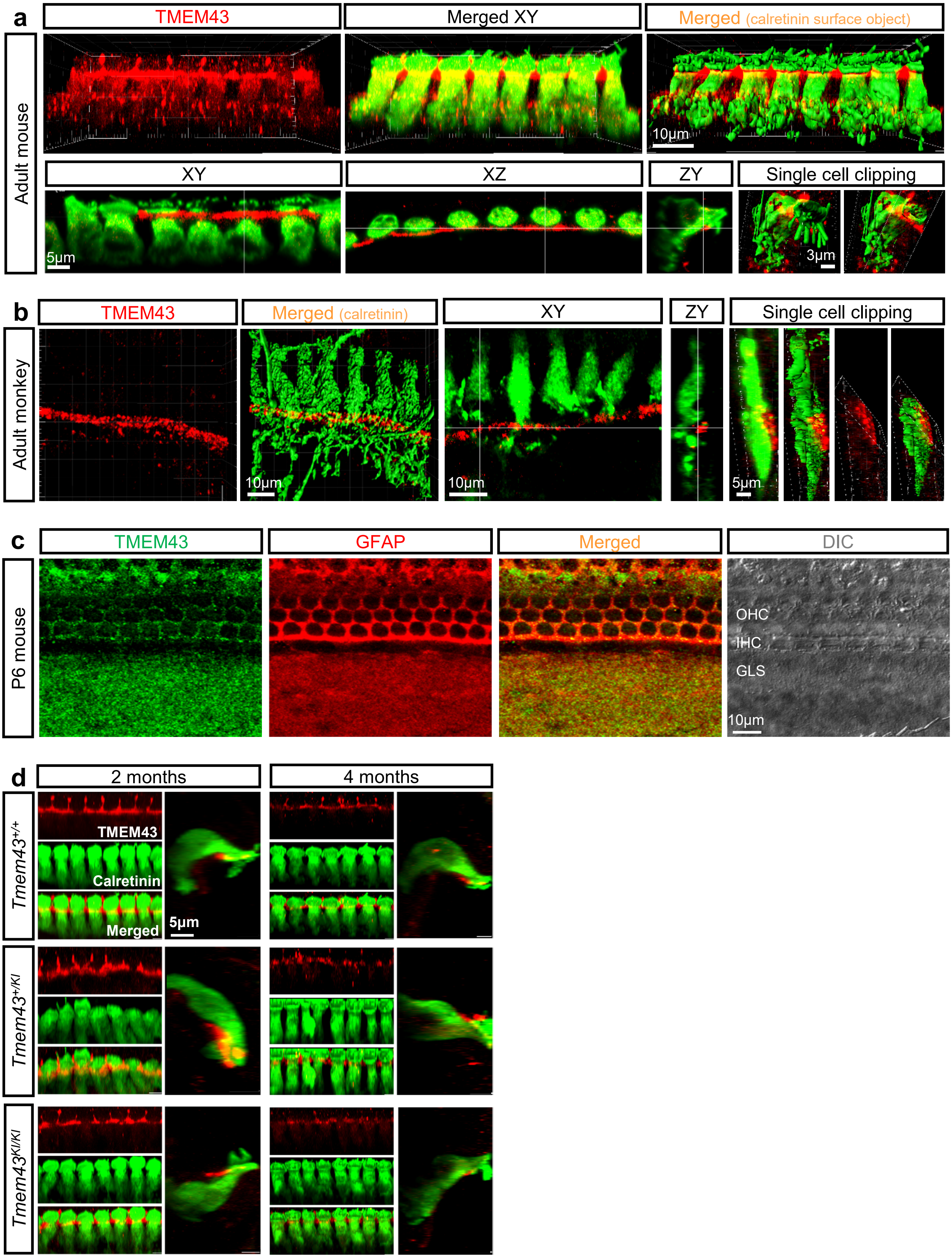

**Extended Data Fig. 3: TMEM43 is highly expressed in cochlear GLSs from developmental stage to adult stage in both mouse and monkey.**

**a, b** Imaris image of cochlea display TMEM43 expression (red) in GLSs that does not co-localize with hair cells (green) in adult mouse (**a**) nor adult monkey (**b**). **c**, Immunohistostained cochlea tissue with TMEM43 and glial marker GFAP. **d**, Confocal micrographs of the organs of Corti from *Tmem43*$^{+/+}$, *Tmem43*$^{+/KI}$ and *Tmem43*$^{KI/KI}$.

**Extended Fig.4**

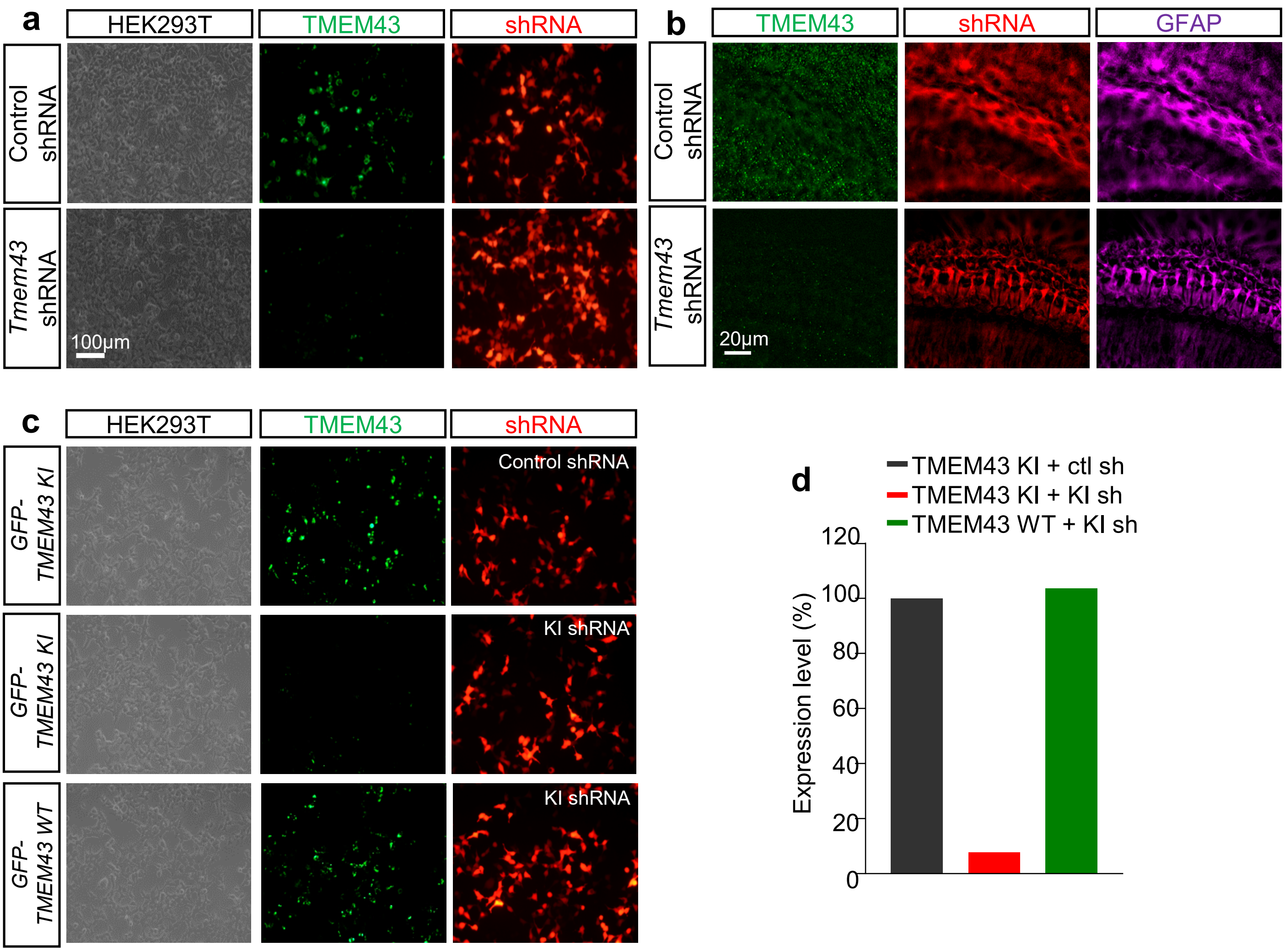


**Extended Data Fig. 4: Knockdown efficiency test of *Tmem43*-targeting shRNA and *Tmem43* KI-sequence targeting shRNA**

**a**, GFP-*Tmem43* transfected with each scrambled and candidate *Tmem43* shRNA on HEK293T. *Tmem43* shRNA suppressed TMEM43 expression by 81.7% (n=20). **b**, Genetic knockdown of TMEM43 with *Tmem43* shRNA virus on cultured mouse cochlea. *Tmem43* shRNA infected cochlea displays reduction in TMEM43 signals at GLSs. **c**, GFP-*Tmem43* or GFP-*Tmem43 KI* was transfected with control shRNA or *Tmem43 KI* shRNA on HEK293T. *Tmem43 KI* shRNA knock-downed TMEM43 KI but not TMEM43 WT. **d**, Quantification of  the protein expression level from (**c**).

### Extended Fig.5

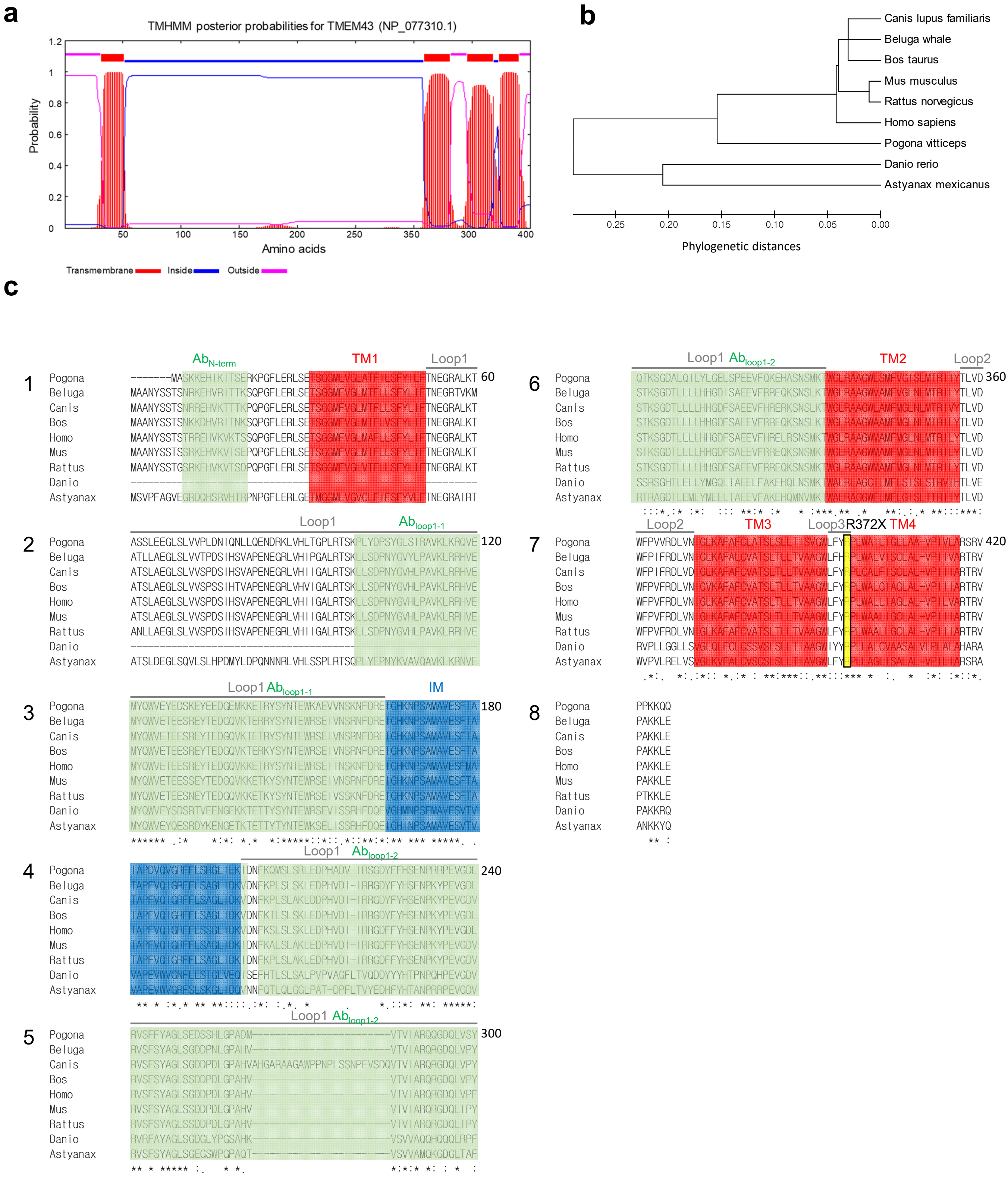

**Extended Data Fig. 5: Phylogenetic analysis of TMEM43 amino acid sequences and structure prediction based on Hydrophobicity analysis.**

**a,** Secondary structure prediction of *Homo sapiens* TMEM43 using 'TMHMM v2.0' server. **b,** The phylogenetic relationship among the 9 different species was inferred using the Neighbor-Joining method by MEGA version 5 software. The optimal tree with the sum of branch length = 1.07711891 is shown. The tree is drawn to scale, with branch lengths in the same units as those of the phylogenetic distances used to infer the phylogenetic tree. **c,** TMEM43 amino acid sequences of *Pogona vitticeps, Beluga whale, Canis lupus familiaris, Bos Taurus, Homo sapiens, Mus musculus, Rattus norvegicus, Danio rerio* and *Astyanax mexicanus* were aligned using 'Clustal Omega' server. The transmembrane (TM) structure with high hydrophobicity for each sequence is represented in red, intramembrane (IM) domain in blue and ANSD causing p.R372X site and C354 are marked with black lining. Epitopes for antibodies (Ab) used in this study are marked with yellow and green.

#### Extended Fig.6

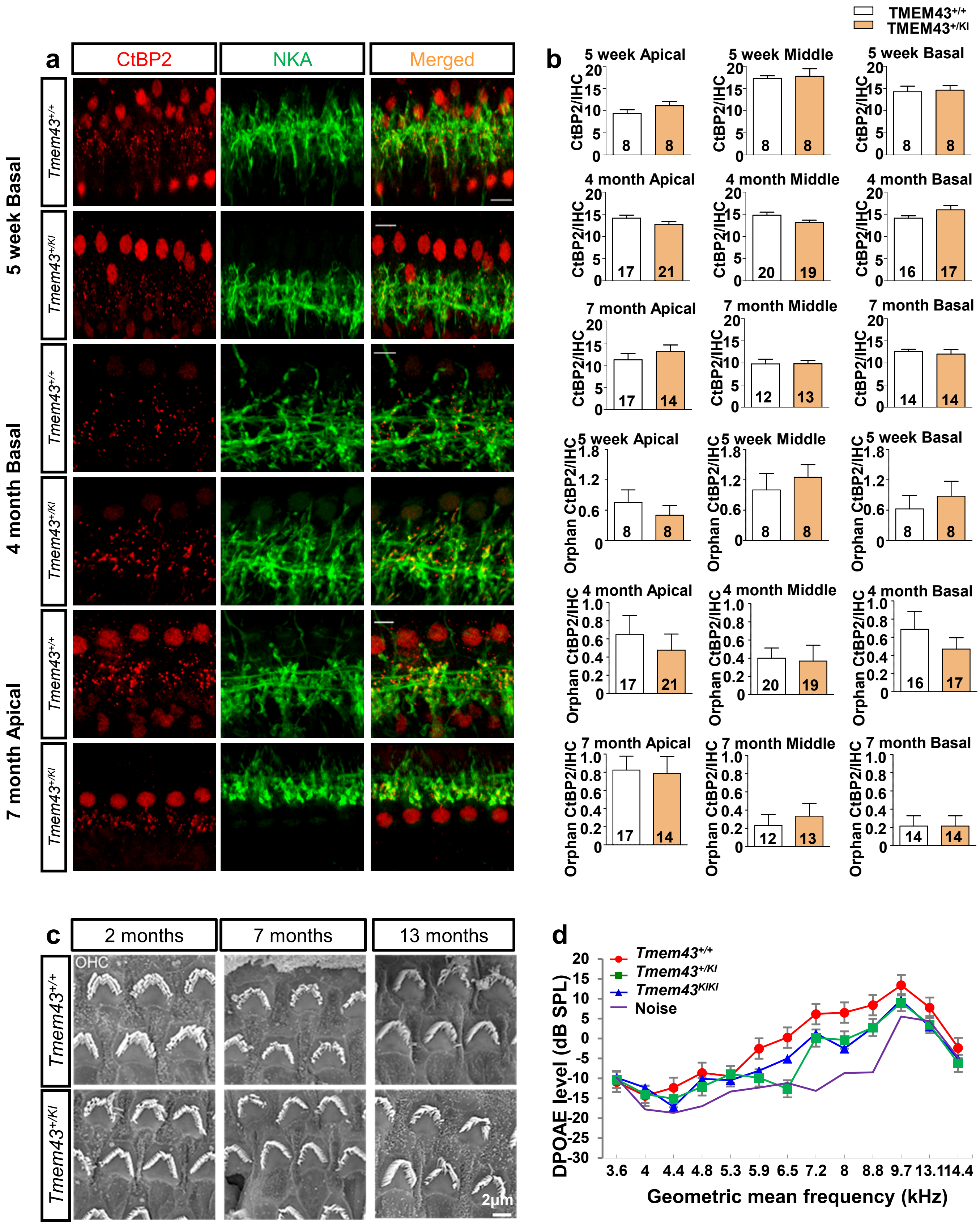

**Extended Data Fig. 6: *Tmem43$^{KI}$* mice display normal synapses and outer hair cell formation.**

**a**, Immunofluorescence staining for CtBP2/RIBEYE and NKA (green) to visualize IHC ribbon synapses in 5-week, 4-month and 7-month-old *Tmem43$^{+/+}$*and *Tmem43$^{+/KI}$* mice. Most CtBP2 puncta juxtapose NKApositive afferent nerve terminals. scale bar = 10 µm, 5 µm, and 5 µm, respectively. **b**, Number of CtBP2 puncta and orphan CtBP2 puncta per IHC (mean ± SEM) in the apical, middle and basal area of cochlea of *Tmem43$^{+/+}$* and *Tmem43$^{+/KI}$* mice. n represents number of IHCs analyzed in each group. Atall ages and cochlear regions examined, there was no statistical difference in the number CtBP2 puncta between genotypes. **c**, SEM showing normal appearance of OHC bundles in 2, 7, and 13-month-old *Tmem43$^{+/KI}$* mice compared to *Tmem43$^{+/+}$* controls. Representative images of whole mounts from apical turns of the cochlea of *Tmem43$^{+/+}$* and *Tmem43$^{+/KI}$* mice. Scale bar: 2µm. **d**, Mean ± SEM growth functions of DPOAE thresholds from 3.6 to 14.4 kHz for littermate *Tmem43$^{+/+}$* (red, n=3), *Tmem43$^{+/KI}$* (green, n=3), and *Tmem43$^{KI/KI}$* (blue, n=3) measured at 6 months after birth. The DPOAE responses from *Tmem43$^{+/KI}$* and *Tmem43$^{KI/KI}$* are not significantly different from *Tmem43$^{+/+}$* (ANOVA, $p > 0.05$) at whole kHz, and significantly higher than noise level (purple) at the frequencies from 7.2 to 9.7 kHz.

**Extended Fig.7**

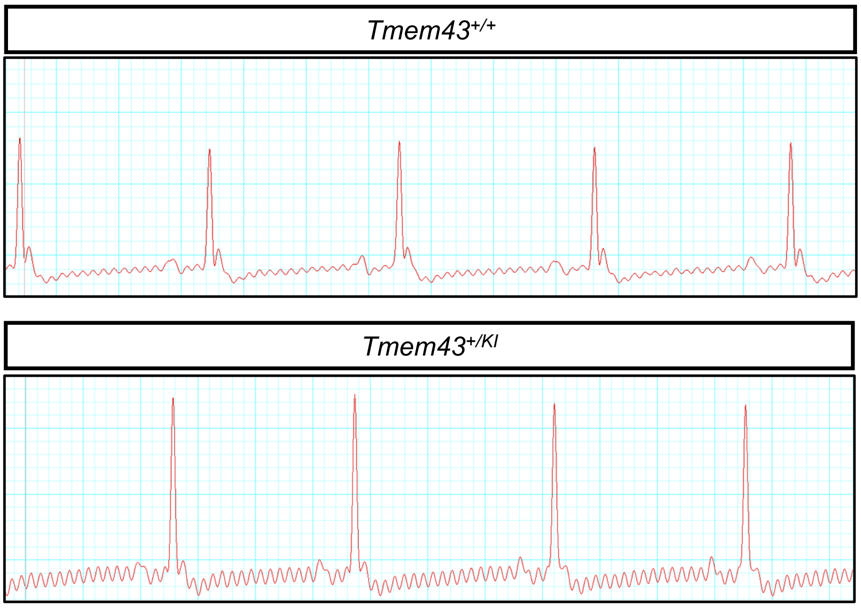


**Extended Data Fig. 7: *Tmem43$^{KI}$* mice have intact electrocardiography**

Representatives of electrocardiography (ECG) from littermate *Tmem43$^{+/+}$* and *Tmem43$^{+/KI}$* measured at 6 months after birth, showing that *Tmem43$^{KI}$* mice display healthy heart function.
