## Supplementary material for "Mutations in TMEM43 cause autosomal dominant auditory neuropathy spectrum disorder via interaction with connexin-mediated passive conductance channels": Suppl Table1

**Supplementary Table 1 | Nonsynonymous variants within the locus[*] co-segregating with ANSD that survived filtering process after exome sequencing among 3 subjects from HN66**

| Func | Gene | ExonicFunc | GenbankID | Position | Exon | Nucleotide | AA | Chr | Start | End | Ref | Alt | Global MAF | KRGDB (N=1722) | PolyPhen-2 | SIFT | GERF++ |
|---|---|---|---|---|---|---|---|---|---|---|---|---|---|---|---|---|---|
| **Exome sequencing** | | | | | | | | | | | | | | | | | |
| exonic | **TMEM43** | stopgain SNV | NM_024334 | Region # 1[**] (3p25.1) | exon12 | c.C1114T | p.R372X | chr3 | 14183206 | 14183206 | C | T | T=0.000008/1 (ExAC) | ND | NA | NA | 4.83 |
| exonic | XIRP1 | nonsynonymous SNV | NM_001198621 | | exon2 | c.T281C | p.M94T | chr3 | 39230656 | 39230656 | A | G | G=0.0000165/2 (ExAC) | ND | 0.998 | 0.784 | 4.93 |
| exonic | VILL | frameshift deletion | NM_015873 | | exon9 | c.946delT | p.F316fs | chr3 | 38040406 | 38040406 | T | - | 0.0001(ExAC) | ND | NA | NA | NA |
| exonic | CTBP2 | nonsynonymous SNV | NM_022802 | | exon5 | c.T2315G | p.L772W | chr10 | 126683123 | 126683123 | A | C | NA | ND | 1 | 0.912 | 5.21 |
| exonic | CHODL | nonsynonymous SNV | NM_ 024944 | | exon5 | c.T665C | p.I222T | chr21 | 19635138 | 19635138 | T | C | C= 0.000008/1 (ExAC) | ND | 0.999 | 0.721 | 5.39 |

[*] Region #1: Chr 3: 13,165,401-22,769,511; Ref: Reference sequence; Alt: Alternate sequence; ND: not detected; NA: not applicable.

The candidate variants were firstly listed up if they were nonsynonymous variants in coding regions. Variants with a minor allele frequency less than 1% were then selected based on the Exome Sequencing Project 6500 (ESP6500), the 1000 Genome Project (1000G), The Exome Aggregation Consortium (ExAC), Trans-Omics for Precision Medicine (TOPMED) and our in-house database containing the exomes of 81 Korean individuals. Following the inheritance pattern, homozygous variants and compound heterozygote variants with sufficient read depths (>10X) and a genotype quality (>20) were selected commonly found in the affected siblings. Finally, the variants which have none of clinical significance of dbSNP ID were finally identified. In silico prediction Algorithm: Polyphen-2 (http://genetics.bwh.harvard.edu/pph2/index.shtml) ; SIFT (http://sift.jcvi.org/www/SIFT_chr_coords_submit.html); Conservation tools: GERP++ score in the UCSC Genome Browser (http://genome-asia.ucsc.edu/) ; ExAC, Exome Aggregation Consortium (http://exac.broadinstitute.org/) ;1000 Genomes (https://www.ncbi.nlm.nih.gov/variation/tools/1000genomes/) ; KRGDB, Korean Reference Genome DB (http://152.99.75.168/KRGDB/); TOPMED, Trans-Omics for Precision Medicine (https://www.nhlbiwgs.org/).
