## Supplementary material for "Mutations in TMEM43 cause autosomal dominant auditory neuropathy spectrum disorder via interaction with connexin-mediated passive conductance channels": Suppl Table2

**Supplementary Table 2.** Non-coding region variants within region#2 that survived filtering process after whole genome sequencing among 8 subjects from SB162

| Chr | Start | End | Ref | Alt | Func.refGene | Gene.refGene | MAF | Affected group | Unaffected group | Normal Control |
|---|---|---|---|---|---|---|---|---|---|---|
| chr3 | 2722187 | 2722187 | G | A | intronic | CNTN4 | T=0.0002/1 (1000 Genomes)<br>A=0.000008/1 (TOPMED)<br>A=0.00003230/1 (gnomAD) | | | ● |
| chr3 | 3449476 | 3449476 | C | T | intergenic | CRBN,LRRN1 | ND | | | ● |
| chr3 | 3630012 | 3630012 | G | A | intergenic | CRBN,LRRN1 | ND | | | ● |
| chr3 | 3765131 | 3765131 | T | - | intergenic | CRBN,LRRN1 | -=0.0766/9615 (TOPMED)<br>-= 0.07394/2093 (gnomAD) | X | | |
| chr3 | 3774063 | 3774063 | T | G | intergenic | CRBN,LRRN1 | G=0.00002/3 (TOPMED)<br>G= 0.00003228/1 (gnomAD) | | | ● |
| chr3 | 4300751 | 4300751 | A | G | intergenic | LRRN1,SETMAR | ND | X | | |
| chr3 | 4352459 | 4352459 | - | A | intronic | SETMAR | A=0.0001/15 (TOPMED) | | | ● |
| chr3 | 4619624 | 4619625 | AC | - | intronic | ITPR1 | -=0.0002259/7 (gnomAD)<br>-=0.0003704/6 (EAS- gnomAD) | | | |
| chr3 | 4619626 | 4619626 | - | TC | intronic | ITPR1 | -=0.0002259/7 (gnomAD)<br>-=0.0003704/6 (EAS- gnomAD) | | | |
| chr3 | 5003582 | 5003582 | A | G | ncRNA_intronic | BHLHE40-AS1 | ND | X | | ● |
| chr3 | 5452014 | 5452014 | A | G | intergenic | MIR4790,GRM7-AS3 | ND | | | ● |
| chr3 | 5700196 | 5700196 | C | T | intergenic | MIR4790,GRM7-AS3 | T=0.00006459/2 (gnomAD)<br>T=0.001250/2 (EAS- gnomAD) | | | |
| chr3 | 12365259 | 12365259 | - | A | intronic | PPARG | ND | | ● | |
| chr3 | 12531584 | 12531584 | A | G | intronic | TSEN2 | ND | | | ● |
| chr3 | 13023564 | 13023564 | C | T | intronic | IQSEC1 | ND | | | ● |
| chr3 | 13229090 | 13229090 | G | A | intergenic | IQSEC1,NUP210 | ND | | | ● |
| chr3 | 13792436 | 13792436 | T | C | intergenic | LINC00620,WNT7A | ND | | | ● |
| chr3 | 14079864 | 14079864 | A | G | ncRNA_intronic | TPRXL | ND | | | ● |
| chr3 | 14183206 | 14183206 | C | T | Exonic_stopgain | TMEM43 | T=0.000008/1 (ExAC)<br>T=0.000008/1 (TOPMED)<br>T=0.00001 (2/246212, GnomAD) | | | |

3 X, variants not detected in affected group; ●, variants detected in unaffected group, normal control
